# *Suregada multiflora* Root Fluro-PolyPhosphate-Glycoside (NU2) inhibits MDR Bacteria and Unicellular Parasites Targeting DNA Topoisomerase I

**DOI:** 10.1101/2024.10.23.619771

**Authors:** Asit Kumar Chakraborty

## Abstract

**Background:** Common antibiotics were useless now as *mdr* genes were gradually accumulating in all environmental bacterial plasmids. Thus, development of alternate to synthetic antibiotics is great.

**Objectives:** Plants secrete ex-metabolites to prevent soil bacteria and thus a good source for new antibiotics. Our objective is to discover an abundant active phyto-chemical against MDR pathogens from household plants of West Bengal.

**Methods:** *Suregada multiflora* plant was grown in clay pot. Root ethanol extract was made for overnight at RT and concentrated. Preparative TLC was performed to isolate major active component by UV-shadow method and was further purified by HPLC. NU2 Inhibitory assays on MDR *Escherichia coli* KT-1, *Leishmania donovani* promastigotes, *Plasmodium falciparum* in human RBC culture were performed as described previously. Biochemical assays, elementary and spectral analyses (MASS, FTIR, NMR) were performed to validate chemical nature of NU2. DNA Topoisomerase I, RNA polymerase and DNA polymerase assays were performed to find modes of action. The structures of NU2 were drawn using Stacher and NU2 binding to DNA topoisomerase I was performed using SwissDock server [Bugnon et al. 2024].

**Results:** Well cultivated *S. multiflora* root extract was 18fold active than bark extract and further enhanced 2.8fold in MS media root culture. The four times TLC purified active principle (NU2) was eluted at 10.2min in HPLC C_18_ column. NU2 inhibits MDR bacteria as well as unicellular parasites *Plasmodium falciparum*, *Leishmania donovani* and Trypanosoma brucei. Spectral analyses suggested NU2 was a glycoside-polyphosphate with molecular mass 454mu and molecular formula C4H7P4O16F. FT-IR gave broad band at 3000-3600cm^-1^ for -OH including at 1559cm-1 for -OH bending. The distinct phosphate -PO4^-3^ group absorption was confirmed at 1021.4cm^-1^ including 928.7cm^-1^ for C-O-P and skeleton minor phosphate group vibration at 525cm-1. Proton-NMR demonstrated huge absorption at δ=2.16ppm for C-O-H, at δ=1.52ppm for -POH while Carbon-NMR at 30.1ppm for C-H and at 207.01ppm for -HC=O. NU2 actively inhibited the DNA topoisomerase I but DNA polymerase and RNA polymerase of *Escherichia coli* were not inhibited. The AI-guided computer simulation suggested that NU2.1, NU2.2 and NU2.3 were better inhibitors for topoIII than topoI of *E. coli*, *M. tuberculosis* and *L. donovani*. We postulate that NU2 prevent the DNA nicking, strand passing and ligase activities after specific binding to the domain-2 active sites.

**Conclusion:** Thus, NU2 cyclic glycoside-fluoropolyphosphate may be a versatile type-I DNA topoisomerase inhibitor and could be studied further for drug design against bacterial and parasite drug-resistant pathogenesis.

**Graphical Abstract:** 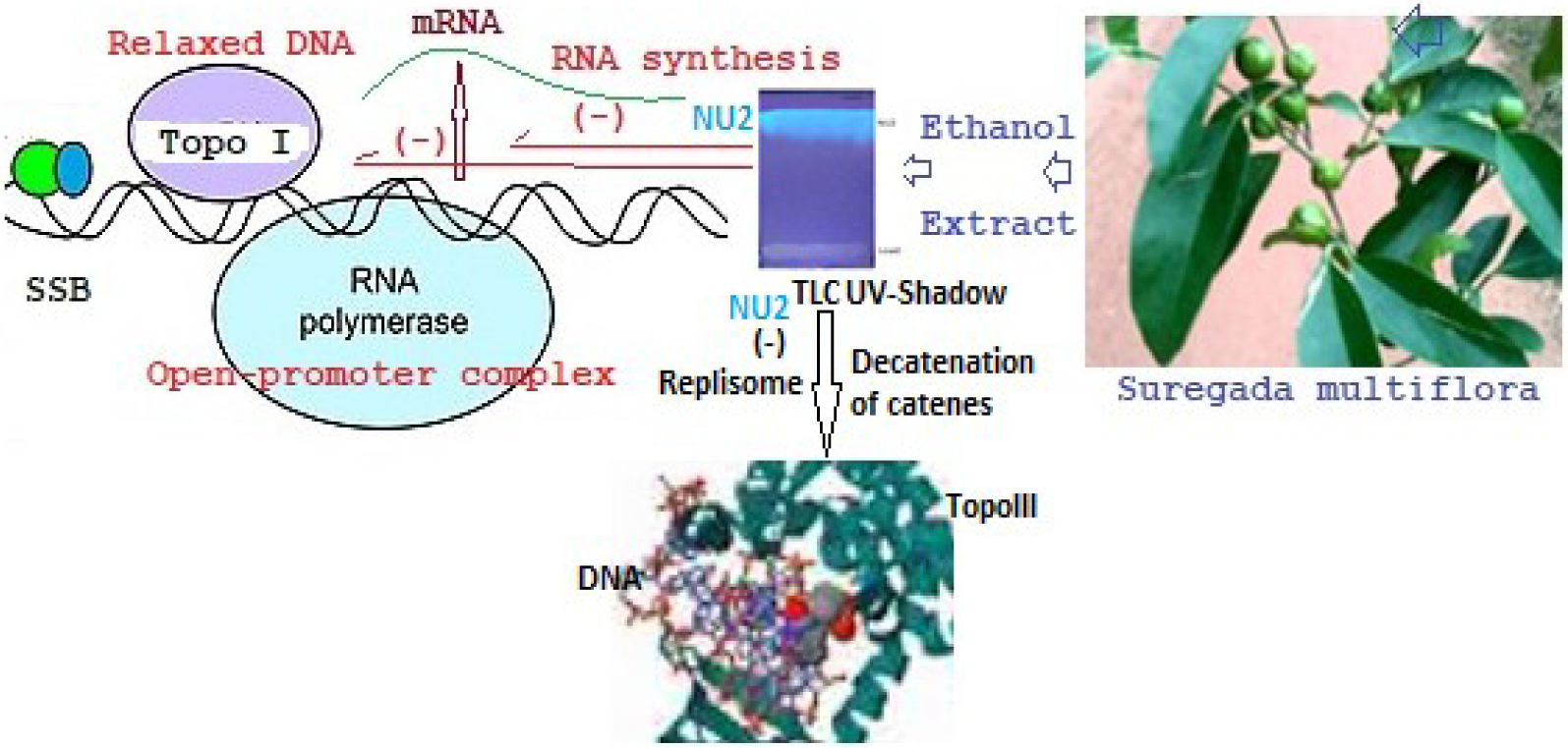

## Introduction

Human suffering from diseases is ancient and universal. Medicine to cure such ailments is epistemologically linked to scientific development and social structure at that time and its prevalent theories may vary time to time [Hong-Fang, et al., 2009]. Indian medicine is devoted to plant remedies and has defined the way of salvation from suffering based on Indian Philosophy build by Patanjali, Kanada, Kapil and Buddha; followed by Ayurveda scientists like Sushruta, Charaka and Chakrapani as depicted in Susruta Samhita, Charaka Samhita and Atharva Veda [Mukherjee, et al., 2017]. During Vedic time (1500BC), it was believed that many rituals of God are central to maintain the order of cosmos and many ceremonies helps the universe to keep working smoothly defeating demons. In Veda, trees are God and each medicinal tree is worshiped traditionally [Touwaide & Appetiti., 2013]. Those things are not useful today as scientific discoveries emerged by Pasteur, Flaming & Washman, Watson & Crick, Khorana, Sanger, Baltimore, and Bishop deciphering the role of invisible microbes in diseases and its cure by antibiotics and phyto-chemicals, DNA structure and its information flow from RNA to protein and how changes in oncogenes (mutation and promoter activation of Ras and Src) cause cancer and gene therapy technologies [Chakraborty, et al., 1991., 1993; Dunbar et al., 2018]. Further defining the atomic model, crystal structure and orbital bonding by NMR, FT-IR and MRI have greatly deepen our understanding of nanostructures in biology and medicine. Organic synthetic chemistry and virtual screening of drugs are so instrumental that computer and robotics have rooted in our mind and such social infrastructure is very ideological to Vedic essential rituals. Still, it appears that we cannot ignore the plant extracts today similar to ancient days as the methods of active ingredients identification by gel chromatography, HPLC and its chemical structure identification (NMR, MASS, FT-IR) are possible quite accurately today [Chakraborty et al., 2022]. Truly, many modern widely used drugs like aspirin, taxol, topotecan, quinine, and artemisinin are of herbal origin but effectively cure bacterial infections, cancer and malaria [Chakraborty., 2019a].

Recently, few thousand tons of antibiotics were made and consumed by 800 million peoples of this Earth every year and we saw contamination of antibiotics in sea and river water fostering drug resistance microbes [Finch., 1998]. WHO warned the unnecessary use of antibiotics in human, animal and agricultural land fostering MDR genesis in the gut and its outbreaks. But the situation now beyond control and >95% of isolated clinical bacteria from human and animal are ampicillin resistance (at 50µg/ml) and also in sea water similar situation with >40% resistance has been reported [Chakraborty., 2015]. However, 3^rd^ generation cephalosporins have 0.2% (at 5µg/ml) and carbapenems have 0.002% (at 1µg/ml) resistance in Ganga River water. Interestingly, the rate of resistance is increasing steadily year to year. As for example, the approximate percentage of increase in *Salmonella typhi* infections between 1987-1994 were reported as: ampicillin 18 % to 78%, chloramphenicol 15% to 78%, tetracycline 53% to 89% and co-trimoxazole 3% to 37%. The ampicillin resistance in *Haemophilus influenzae* isolates were increased from 4% in 1977, to 31% in 1981 and now is almost 100%. TB anti-microbial resistance has reached astonishingly high in India with 20 million MDR infections of *Mycobacterium bacilli* that has been treated by rifampicin and streptomycin drugs in the past. The *RpoB*, *pncA*, *ponA, penA,* and *rpsL* genes mutations are involved in multi-resistance in TB and Gonorrhoea [McArthur, et al., 2013]. Multidrug resistant STDs are increasing in the East and Western countries at the same level as many chromosomal membrane-bound drug efflux proteins (macA/B, mtrCDE, mexAB/CD/EF, emrA/B-TolC) are activated [Chakraborty., 2019b]. In one ward, large (100-500kb) MDR conjugative plasmids carry 5-15 *mdr* genes, 4-16 *Tra* genes, 4-24 transposases, integrases and recombinases, few MFS/RND transporters, few transcription factors and more sadly 10-30 unknown genes. Our study with Ganga River water has demonstrated that super-drugs like imipenem, colistin, amikacin, linezolid, lomofloxacin resistant species are increasing in the environment. No antibiotic is available to treat NDM-1 *Escherichia coli*, XDR *Pseudomonas aeruginosa*, MRSA *Staphylococcus aureus* and MDR *Acinetobacter baumannii* as *blaNDM-1, mcr-1, aac3’-1b-cr, strA/B,* and *sul1/2/3* genes were accumulated in those plasmids with target specific mutations in *gyrA* (fluroquinolone Resistance), *rRNA* (aminoglycosides resistance) and *porB* (cephalosporins resistance) genes [Chakraborty et al., 2017a; 2019c].

Release of heavy metals in rivers is huge due to mineral extraction and data indicated so many metal efflux gens and metal detoxification genes in MDR plasmids (Chakraborty & Roy., 2020]. Plants release ex-metabolites in defence of soil and water bacteria. Thus, we thought that plants were a unique source of antibiotics to control MDR bacteria. To solve unstable nature of phytochemicals we thought three possible remedies: (a) search for more active and stable phyto-extracts, (b) cultivation of plant in big clay pot keeping in local urban area, and (c) plant tissue culture. Here, we showed that *Suregada multiflora* root and bark derived NU2 glycoside-polyphosphate was an excellent antibiotic to kill MDR bacteria as well as single-cellular parasites like *Leishmania donovani*, *Trypanosoma brucei* and *Plasmodium falciparum*.

The drug was targeted type I DNA topoisomerases and crystal structures of such enzymes were known to perform docking analysis [Patel et al. 2010; Baker et al. 2009; Perry and Mondragón, 2003; Chen and Wang, 1998]. A single copy top1A gene (protein id. AEJ52168; 5D5H_A) was located in *Mycobacterium tuberculosis* and quite similar to *Escherichia coli* enzyme (protein id. 1I7D_A) [Tan et al. 2016]. Genetic analyses identify *LdTOPIA* gene located on *L. donovani* chromosome 34, encoding for a 636-amino acid polypeptide (Protein id. AAG27713; AAD41814) while topoIII (867aa) was located in chromosome 28 (protein id. CAJ1990333). The *Plasmodium* Type-I topoisomerase is a monomeric protein of 839 amino acids (protein id. CAA58716; XP_001351663) while topo3 is 710aa (protein id. XM_001350149) [Bansod et al. 2022]. The *PfTopI* gene appears as a single copy on chromosome 5 of the plasmodial genome. Structurally, the enzyme well resembles other topoisomerases, keeping a 42% homology with the human enzymes (protein id. ADP90493, AAA61207, 1T8I_A). We firmly studied the topoisomerase-NU2 binding using SwissDock Server (Röhrig et al. 2023) and believed a better inhibitor for DNA topoisomerase III (protein id. BAA15551; CAJ1990333; 1I7D_A). This is the first report that *Suregada multiflora* has abundant cyclic fluoro-polyphosphate which is an inhibitor of type-1 DNA topoisomerases.

## Materials & Methods

### Isolation and characterization of MDR bacteria from Ganga River

Water from Ganga River was collected in the morning from Babu Ghat (Kolkata 700001) and Howrah Station area (Howrah 711101). About 100µl of water was spread onto 1.5% Luria Bartoni-agar plate containing different antibiotics at 5-50µg/ml. MDR bacteria were selected in agar-plate containing ampicillin, streptomycin, chloramphenicol, tetracycline or ciprofloxacin at 50, 50, 34, 20 µg/ml respectively [Chakraborty., 2015]. Antibiotics susceptibility test was conducted for determine the sensitivity by Kirby-Bauer method. The results were determined by criteria from Clinical and Laboratory Standards Institute (CLSI) [Ausubel et al., 1989].

### Culture of malaria parasites and assays of drug inhibition

Chloroquine sensitive *Plasmodium falciparum* (strain NF54) was cultivated at 37^0^C, in O+ erythrocytes under 3% O_2_, 4% CO_2_, 93% N_2_. Parasitized 100µl RBC cells (0.25% parasitemia) were taken into wells of microtiter plate and equal volume of OPTI-M1640 (GIBCO-BRL) medium containing 20% heat inactivated serum added and incubated for 48hrs. in presence or absence of drug. After 48hrs 25µl medium containing 2.8 µCi of ^3^H-Hypoxanthine (DuPont, NEN, 2.8mCi/ml in 50% ethanol) was added to each well and incubated another 24hrs. Total cells were harvested into GF/C filter papers using Cell Harvester and counted into a Scintillation counter [Meyer., et al., 2018].

### MTT assay of Leishmania donovani

The effect of phyto-extracts on the viability of *L. donovani* AG83 was determined by 3-(4,5-dimethylthiazol-2-yl)-2,5-diphenylterazolium bromide (MTT) assay [Kumar, et al., 2018]. The cells at the exponential phase were collected and transferred into 24-well plate (approximately 4×10^6^ cells/well). The cells were then incubated for 12hrs in the presence of various concentrations of the extracts. After incubation, the cells were centrifuged, and then finally suspended in 100μl of PBS in 96-well plates. Ten microliters of MTT solution (10μg/ml) were added in each sample well plates, and the samples were incubated for 4 hrs. After incubation, 100μl of stock solution (stock: 496μl of isopropanol and 17μl of cons. HCl) was added and kept for 20min at room temperature. The optical density was taken at A570 on an ELISA reader (Multiscan EX, Thermo Fisher Scientific, Waltham, MA).

### Growth of molly fishes

Molly fishes (Black in colour, 1.2cmx3mm) were cultured in 1L jar with Kolkata Municipal Corporation water (500ml, deionised) supplemented with grinded rice, wheat or commercial pellet as food source. The growth and movement were observed by naked eye and if the fishes died (floats) within 1-3 days was taken as potential toxicity as compared to standard. **Growth of HeLa cells**

HeLa cells were grown in Dulbecco Minimal Eagle Medium (6cm plates, 3ml media, 10^5^ trypsinized cells) containing 10% foetal bovine serum at 37°C and 5% CO_2_ as described previously. Contact inhibited mono-layer cells were observed within 72hrs and picture was taken at 400X. If the confluence was not reached, then it was taken as growth rate inhibition by drug at that concentration. The plant extract treated (72hrs) HeLa cells were divided into fresh media plates at 1:8 and if the cells did not grow at all was taken as potential toxicity [Chakraborty, et al., 1995].

### Preparation of MS Media and root culture

The plant tissue culture was initiated with MS Media (Murashige & Skoog, 1962). The 20X Macro Elements was made by adding 3.8g KNO_3_, 3.3g NH_4_NO_3_, 0.88 g CaCl_2_,2H_2_O, 0.74g Mg SO_4_, 0.34g KH_2_PO_4_ in 100 ml Water. 1000X Micro Elements was made by adding 2.23g MnSO_4_, 0.86g ZnSO_4_.7H_2_O, 0.62g H_3_BO_3_, 25mg Na_2_MoO_4_.2H_2_O, 2.5 mg CuSO_4_.5H_2_O, 2.5 mg CoCl_3_.6H_2_O in 100ml water. 200xFe+2-EDTA solution was made by adding 745 mg Na_2_EDTA in 75 ml boiling water and gradually adding 557mg FeSO_4_.7H_2_O and volume adjusted to 100 ml with water and kept in Amber colour bottle. 1000xVitamins was made by adding 25mg Nicotinic acid, 25mg Pyridoxine.HCl, 5mg Thiamine.HCl in 50 ml water and store at 0°C for 15 days only. 2000x KI was prepared by adding 83mg KI in 50 ml water and store at 10-16°C. 500X Myoinisitol was made by adding 1g myo-inisitol in 20 ml dH_2_O. Sucrose added at 30-50g/L and glycine was added at 2mg/L. Auxins were prepared by adding 10 mg each 2,4 dichloro phenoxy acetic acid (2,4-D), Indole acetic acid (IAA), α-Napthalene acetic acid (NAA) and 3-indole butyric acid (IBA) in each 1 ml ethanol and adding 9 ml water and store at 0°C. Cytokinines each Kinetin (6-furfural amino purine), BAP (6-Benzyl amino purine), Zeatin, 2iPA (2-isopentenyl adenine) were prepared by adding each10 mg in 1 ml 1N HCl and then add 9 ml water. Final media was prepared by dissolving 7g phyto-agar in 740 ml boiling water and further autoclaving at 15psi/15min-cooled and all ingredients added. Final Mixture of 1000ml MS Media was made by adding 20x Macro = 50 ml, 1000x Micro = 1 ml, 2000x KI-0.5 ml, 200x Fe-EDTA = 5 ml, 1000x Vitamins = 1ml, 1000x glycine = 1 ml, 500x Myo-inositol = 2ml, 7g phyto-agar in 940 ml water.

Roots were collected in the morning and placed in a 500ml beaker covered with cheese cloth and washed with running tap water for 15 min and washed in 3% Tween-20 for another 15 min in a magnetic stirrer, and further treated with 1% HgCl_2_ for 2 min and washed repeatedly with autoclaved water. Roots were incubated in MS media for 5 days and extracted in ethanol for overnight at room temperature.

### Isolation of NU2 phytochemical from Suregada multiflora root ethanol extract

MDR-Cure extract was spotted on the TLC plate and ascending chromatography was performed in 60% methanol+10% acetic acid+30% water as mobile phase. Alkaloid was spotted by UV shadowing and anthroquinones were spotted by spraying the TLC plate with alcoholic KOH (4.5gm KOH dissolved in 20 ml water and mixed with 80 ml methanol). The alkaloid spot was found just above the anthroquinones spot (brown colour). UV shadow technique was used to collect phytochemical band that was excised from TLC plate (silica) and dissolved in ethanol, clarified at 10,000 rpm/5min and used as source of antibacterial phytochemicals. Four times TLC purified product was used for Mass, IR and NMR studies (Chakraborty et al., 2022).

### High Performance Liquid Chromatography

TLC-purified active sample (5mg) was dissolved in 0.5ml methanol, filtered through a membrane filter and 0.1ml sample was loaded onto a HPLC C-18 column equilibrated with methanol. Fractions of 0.5ml were collected and major active peak (retention time-3min) was collected and vacuum dried (Naidoo et al., 2018).

### Phyto-chemicals assays

TLC-purified chemicals were used for biochemical assays. TLC-purified CU1 chemical was dissolved in water and if ethanol extract was used then ethanol concentration was lowered by adding water during assay. *Assay of Saponins*: 0.5 ml plant extract + 2ml methanol + 2ml water. Formation of persistent foam on the surface was taken as an indication for saponin. *Assay of Flavones*: Alkaline Reagent Test: 2 ml of 2% NaOH solution was mixed with 0.4ml plant extract. A concentrated yellow colour was produced, which became colourless with addition of diluted HCl to mixture. *Test for glycosides*: To 1ml plant extract, 3 ml of glacial acetic acid followed by addition of few drops of FeCl_3_ solution were added. Then, 2 ml of conc. H_2_SO_4_ was added giving a reddish-brown colour. *Test for Terpenoids*: 0.5ml chloroform + 0.5 ml sulfuric acid concentrated + 0,5 ml plant extract + 0.5 ml water. Boiled the mixture. A grey colour indicates triterpenes. *Test for steroids*: 0.4ml chloroform + 1ml plant extract + few drops concentrated sulphuric acid. Red colour indicates steroids. *Test for alkaloids*: Alcohol extracts were diluted with dilute HCl (1:5v/v) and treated with few drops of Mayer’s reagent. A reddish brown or orange precipitate indicates the presence of alkaloids [Harborne et al., 1999; Sambrook et al., 1989].

### Fourier Transformed Infra- Red Spectroscopy

HPLC-purified dry active chemical 5mg was mixed with 200mg IR-grade KBr and the tablet was prepared at 13mm Die SET (Kimaya Engineers) at 10Kg/cm^2^. Spectra were taken with a Perkin Elmer Spectrum 100 FT-IR Spectrometer (Serial no. 80944) for 10 min (Griffiths, & de Hasseth., 2007).

### NMR Spectrometry

Proton-NMR spectrometry was performed in CDCl_3_, D_2_O and CD_3_OD (5mg/ml NU2) for 10min. Carbon-NMR was performed in D_2_O only (10mg/ml NU2) in 500MHz JNM-ECX500 Spectrometer (Chakraborty., 2019).

### Assay of *E. coli* DNA topoisomerase I

DNA topoisomerase I was assayed by the reduced mobility of relaxed DNA over supercoiled DNA in a 1% agarose gel in 1xTAE buffer at 15V for 8hrs. The DNA topoisomerase I can nick on one strand and swivels the other strand and then seals the nicks causing CCC-plasmid DNA. In presence of 0.05µg/ml ethidium bromide in gel, such relaxed plasmid DNAs got positively supercoiled and if run for long time, moved faster than negatively supercoiled plasmids. The assay buffer contained 10mM Tris.HCl, PH 7.5, 75mM KCl, 0.5mM EDTA, 10mM MgCl_2_, 0.5mM DTT, 10% Glycerol and 0.1mM PMSF. In a Rx 20µl, then 0.5µg supercoiled plasmid DNA and 1 Unit DNA topoisomerase (New England Biolab) was added and incubated for 30min at 30^0^C and then run 1% agarose gel without Ethidium bromide. After the gel run the, gel was stained with 0.5µg/ml ethidium bromide and take a picture under UV illumination (Chakraborty & Majumder, 1988).

### Assay of Escherichia coli RNA Polymerase using ^3^H-UTP

The assay used to measure RNAP activity was performed as described by Lowe et al (1979) with minor modification [Lowe, et al., 1977; Chakraborty, & Hodgson, 1998]. The reaction (Rx 40µl) was performed with transcription buffer (40 mM Tris-HCl pH 7.9, 200 mM NaCl, 10 mM MgCl_2_, 0.1mM EDTA, 14 mM β-ME, 200nM each ATP, GTP and CTP. 50µM UTP, 2µCi ^3^H-UTP (BRIT, Hyderabad, India), 1.5µg calf thymus DNA and 1U RNA polymerase at 37°C for 20 min. The reaction mixture was spotted onto DEAE-paper pre-socked with 5mM EDTA. The filter paper was then washed with 5% di-sodium hydrogen phosphate, thrice with water and finally with 95% ethanol and dried. The filters were placed into scintillation vials containing 10 ml toluene-based scintillation fluid and counts were recorded on a Tri-CARB 2900TR scintillation counter. Data was normalized and plotted. Rifampicin (10mg/ml in basic ethanol) was used as standard drug and 11 phyto-chemicals (4 times TLC purified; ∼10mg/ml in DMSO) were used [Chakraborty, et al., 2022].

### SwissDock Modelling

The structures were drawn in Swiss-Dock Sketcher and SMILES were formulated in the server [Bugnon et al. 2024]. The *Escherichia coli* DNA topoisomerase I (PDB ID: 1mw8) and DNA topoisomerase III (PDB ID: 1d6M) as well as *Mycobacterium tuberculosis* DNA topoisomerase I (PDB ID:8tfg) were used. The probable structures of postulated NU2 drug (NU2.1, NU2.2 and NU2.3) for docking were presented.

## Result

### Organic phytoextracts from *Suregada multiflora* root and bark are good anti-bacterial

We tested more than 80 plant extracts and obtained five such extracts that inhibited MDR bacteria efficiently. But *Suregada multiflora* and *Cassia fistula* had abundant phytochemical for biochemical and spectroscopic studies and commercialization as drug. Figure-1A showed a *Suregada multiflora* 5 years old plant in clay pot and its abundant secondary roots un-purified (figure-1B) and purified (figure-2C) used to extract phytochemicals in ethanol. Five times concentrated extract or dried solubilised extract (10mg/ml in DMSO) showed antibacterial activity with 15-20mm lysis zone during Kirby-Bauer assay as compared to 20µl of 34mg/ml chloramphenicol or 20µl of 1mg/ml meropenem (20-25mm) using *Escherichia coli* KT-1_mdr (accession no. KY769881) or *Pseudomonas aeruginosa* DB-1_mdr (accession no. KY769875) (see, figure-1). We searched local area of Midnapore (22.658 Lattitude x 87.593 Longitude) and selected *Cassia fistula* (phyto-chemical named CU), *Suregada multiflora* (local name *Narenga*; phyto-chemical named NU) and *Jatropha gossypifolia* (local name *Varenda*; phytochemical named VU) as valuable sources for new drugs to cure superbug infection where no antibiotics work. Figure-2D demonstrated the high content of glycoside-polyphosphate phytochemicals in TLC plate under UV light illumination.

**Fig.1ABC.**
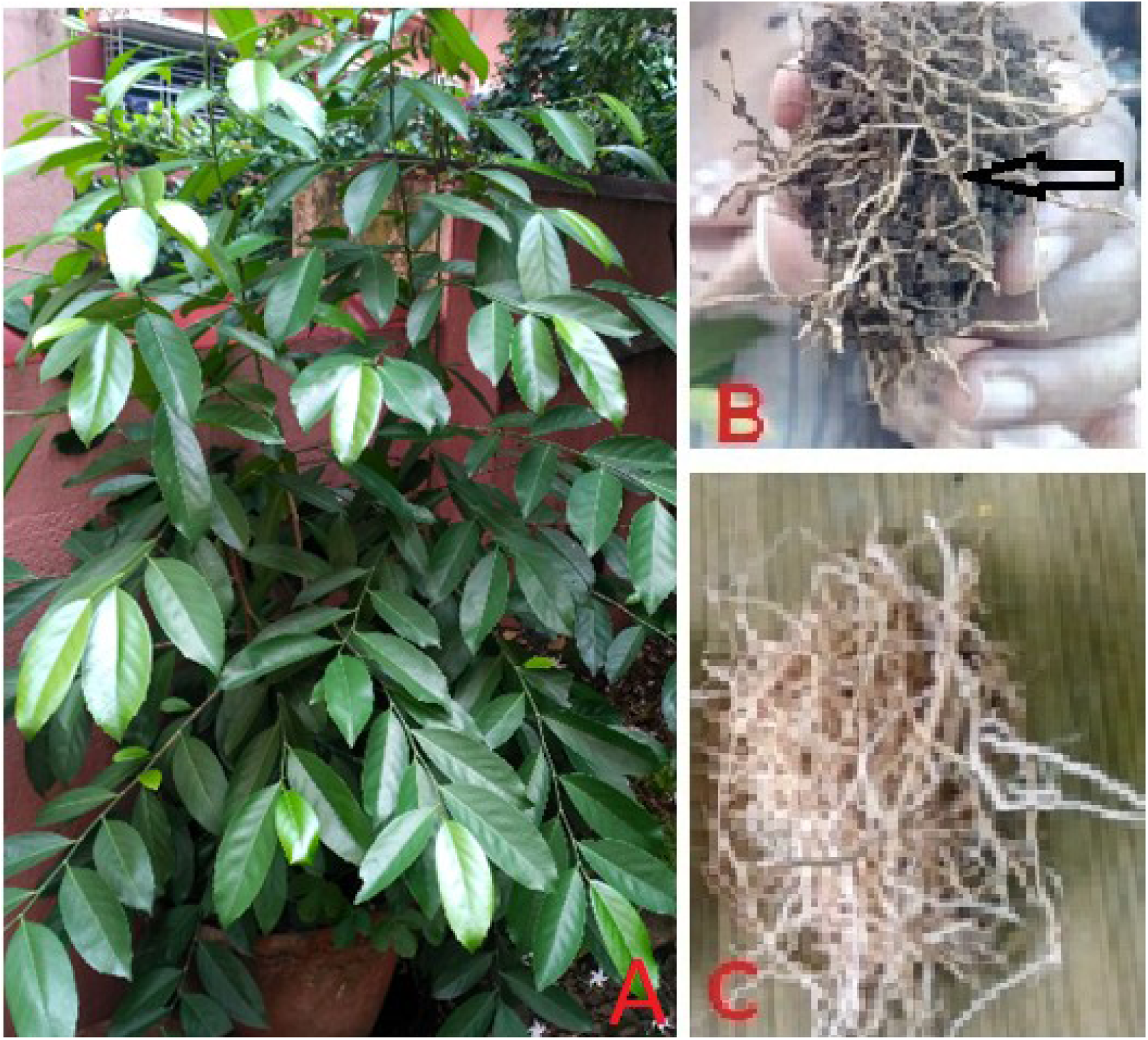
Growth of Naringa plant (*Suregada multiflora*) in clay pot at Kolkata and isolation of roots. Phyto-chemicals are labile on storage. So, plants must at nearby of user. Extracts will be made and consumed very fast to reduce oxidation by air as well as to reduce light induced damage.

**Fig.1D.**
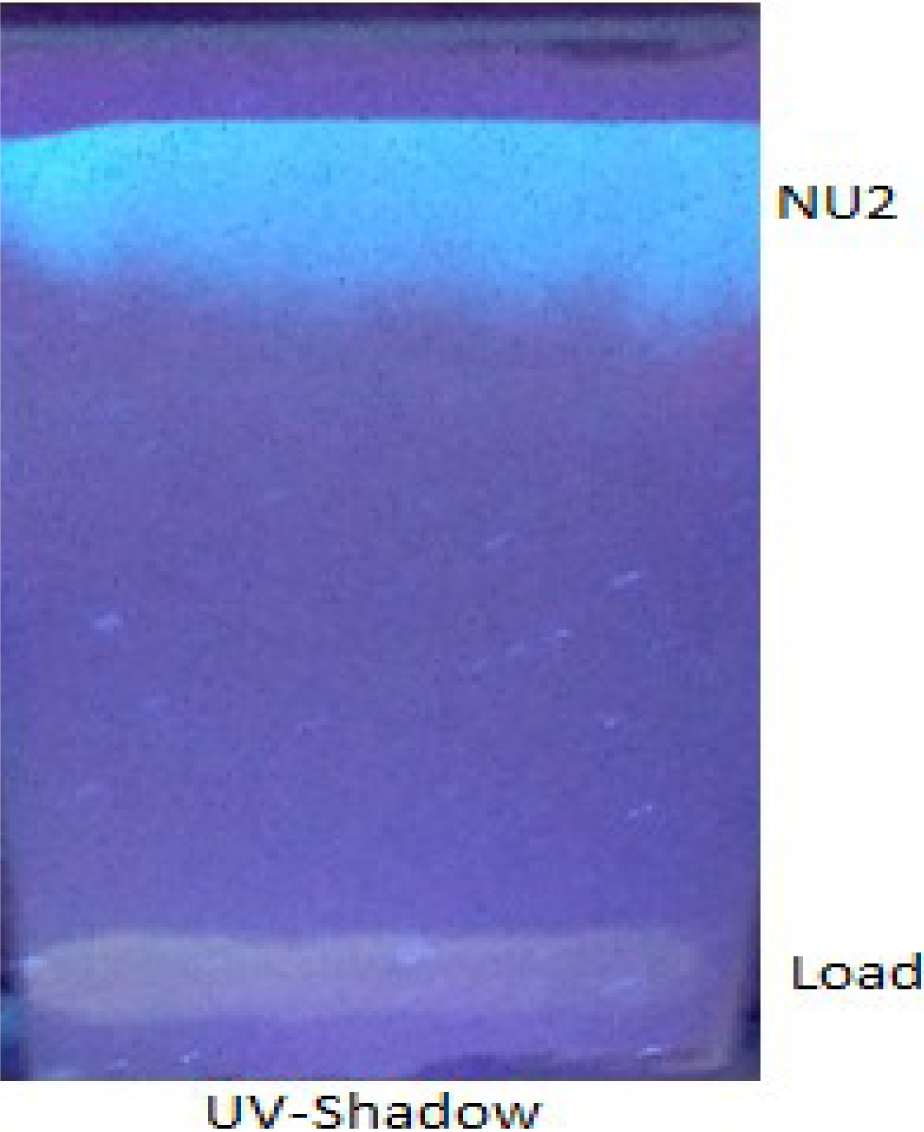
Isolation of NU2 (naringinphos) by repeated TLC and visualization of polyphosphate-glycoside phytochemical by UV-shadow. In this 2^nd^ time TLC of NU2 which ran fast as much purified and concentrated. Otherwise during 1^st^ run preparative TLC the band was found in the middle of the alumina gel.

Figure-2A showed the drug sensitivity assays of organic root extract of *Suregada multiflora* on sensitive bacteria. It appeared *Cassia fistula* bark as well as *Jatropha gossipifolia* root extracts contained good antibacterial compounds that inhibited *E. coli* (plate-1), *B. subtilis* (plate-2) and *S. aureus* (plate-3) whereas last two bacteria were less sensitive to *Cassia fistula* bark extract. Fig-2Bi demonstrated antibacterial activities of *Suregada multiflora* phytochemicals (lane 1) as compared to *Jatropha gossypifolia* root (lane 4) and *Cassia fistula* bark (lane 2). Lane 3 was 20µl of 50µg/ml ampicillin. Figure-2Bii showed the NU2 inhibition of *E. coli*-KT1_mdr in LB media as compared to potent imipenem drug.

**Fig.2A.**
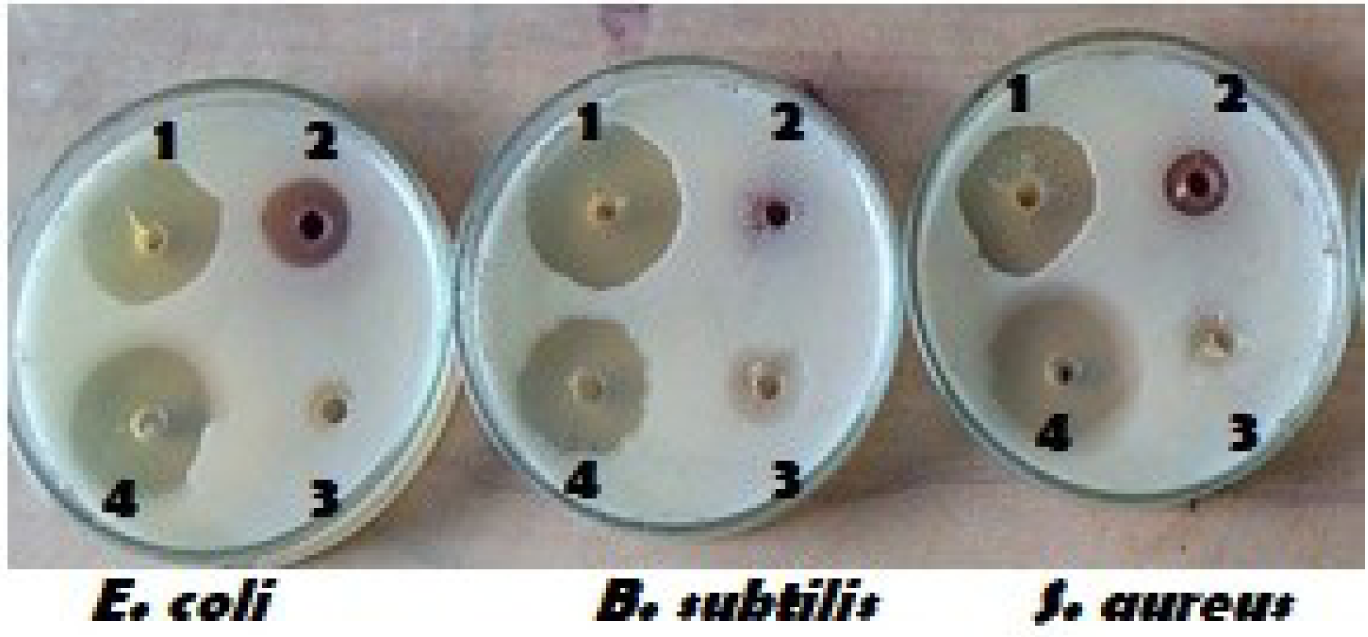
Drug sensitivities of phyto-extracts against sensitive *Escherichia coli, Bacillus subtilis* and *Staphylococcus aureus.* It appeared *E coli* bacteria is more sensitive to *Cassia fistula* bark extract (lane 2) while *Suregada multiflora* extract was equally inhibited three bacteria (lane 1).

**Fig.2B.**
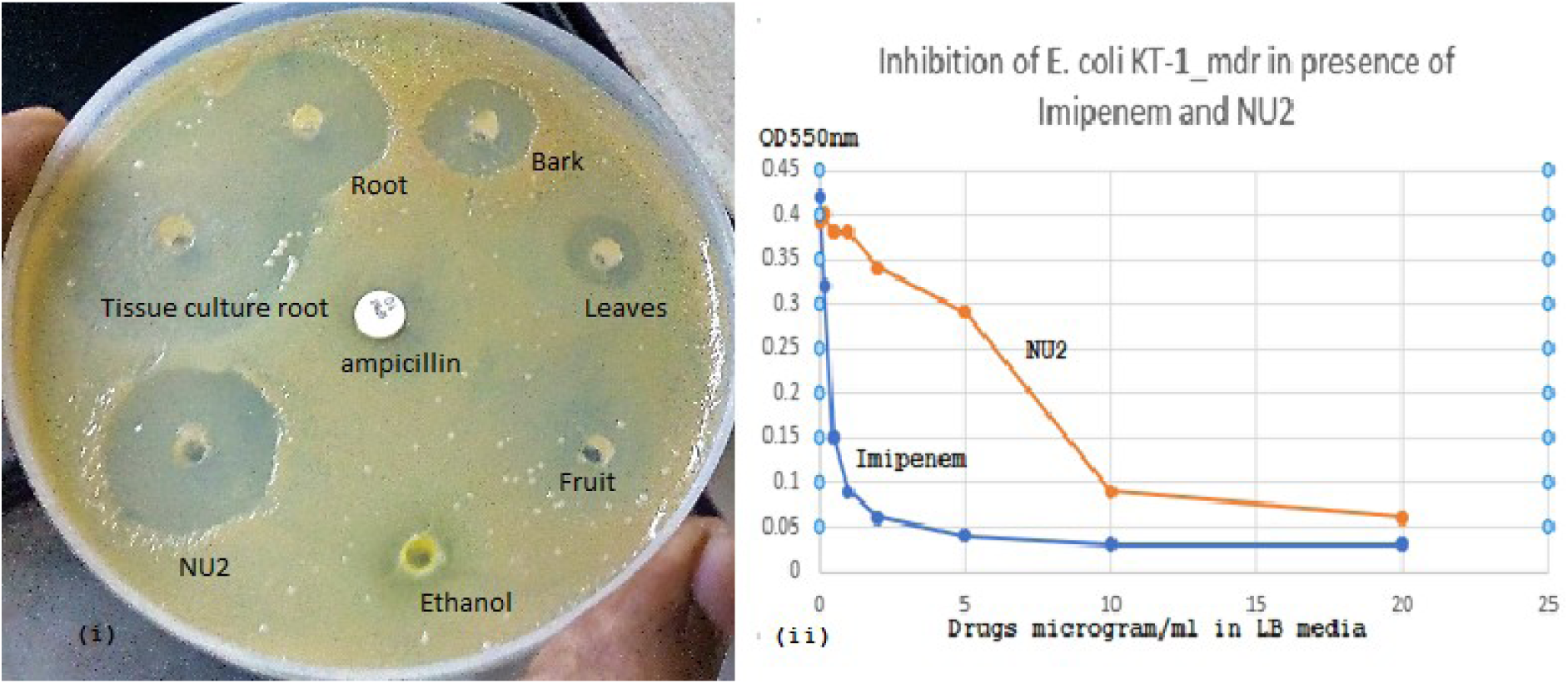
Inhibition of *E. coli* KT-1 superbug by NU2 phyto-chemical. (i) comparative inhibition by organic extracts of *Suregada multiflora* parts, root tissue culture extracts and four times TLC purified NU2. (ii) NU2 in vitro inhibition using LB media as compared to Imipenem, a highly active 5^th^ generation penicillin drug.

### NU2 as traditional anti-parasitic drug against malaria

We checked the anti-parasitic activities against *Plasmodium falciparum*. Suregada multiflora leaves (local name in Bengal: Narenga) were used in the past (before year 1900) to cure malaria (local name: Pali-Azar). I sent some leaves extract to Johns Hopkins and Dr Shapiro showed the extract inhibited *P. falciparum* in vitro. Fig.3A showed the fluorometric detection of *Plasmodium falciparum* inside the human RBC which was inhibited at different concentration of NU2. When we performed radioactive assay using tritiated-Hypoxanthine, the CPM counts reduced from 90,000cpm to 4000cpm in presence of NU2 at 20µg/ml. Similarly, *L. donovani* promastigotes were inhibited at different concentration of NU2 as judged by MTT assays (figure-3B). *T. brucei* blood forms were also lysed at different doses of NU2 as judged by confocal microscope technology (data not shown).

**Fig.3A.**
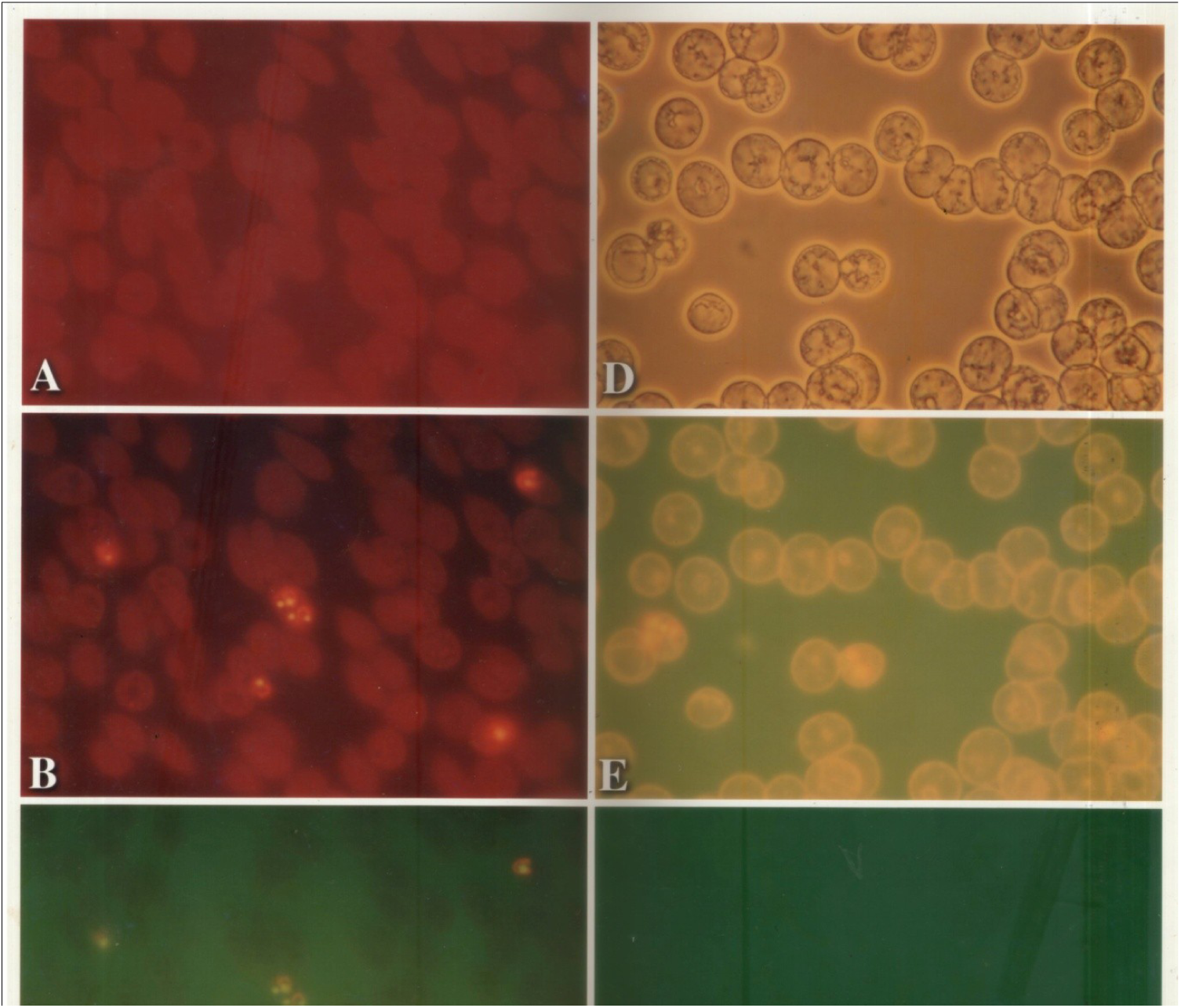
Inhibition of *Plasmodium falciparum* parasites in human RBC by naringinphos. Panel A shows fluorescence of acridine orange treated human pure O^+^ RBC without parasite, Panel B with parasite (red filter), Panel C same as B (green filter), Panel D with parasite (phase contrast picture), Panel E same as D with fluorescence (yellow filter, intense light) and Panel F with parasite + concentrated NU2 (green filter). At 20µg/ml NU2, no parasite was seen. The ^3^H-hypoxanthine labelled normal counts were reduced from 90,000CPM to 4000CPM on NU2 treatment.

**Fig.3B.**
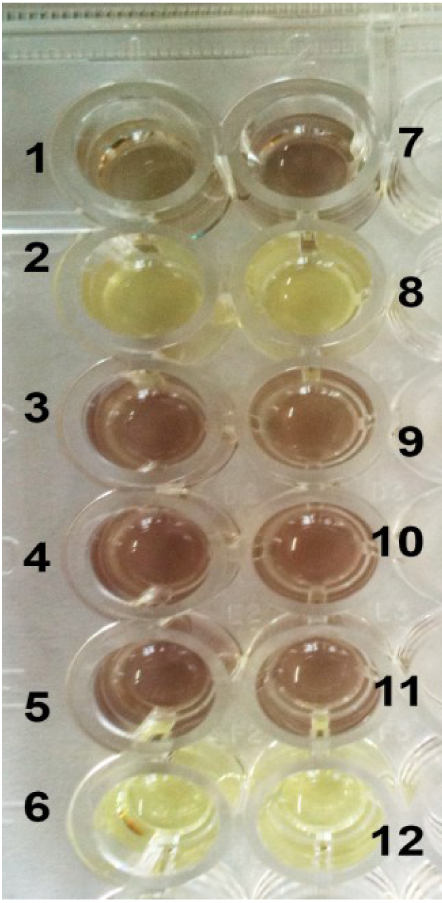
Inhibition of *Leishmania donovani* promastigotes by NU2 *in vitro.* The effect of phyto-extracts on the viability of *L. donovani* promastigotes was determined by MTT assay as described in Material & Methods. Wells 1, 2, 8 were 2, 5 and 10 µg/ml NU2 and wells 3, 4 and 9 were 2, 5, and 10µg/ml CU1, well 7, 10 3% Ethanol control, Wells 5, 11 were untreated cells and wells 6, 12 were reagent blank. It was found that NU2 inhibited the parasite dose-dependent manner.

### Purification of NU2 in HPLC C_18_ column

The NU2 compound was eluted in C_18_ column at 10.2 min and graph predicted there was no much impurities (figure-4). That way our TLC purification was good and the broad band after four successive TLC ran fast in the ascending TLC in 40% methanol plus 10% acetic acid medium.

**Fig.4A.**
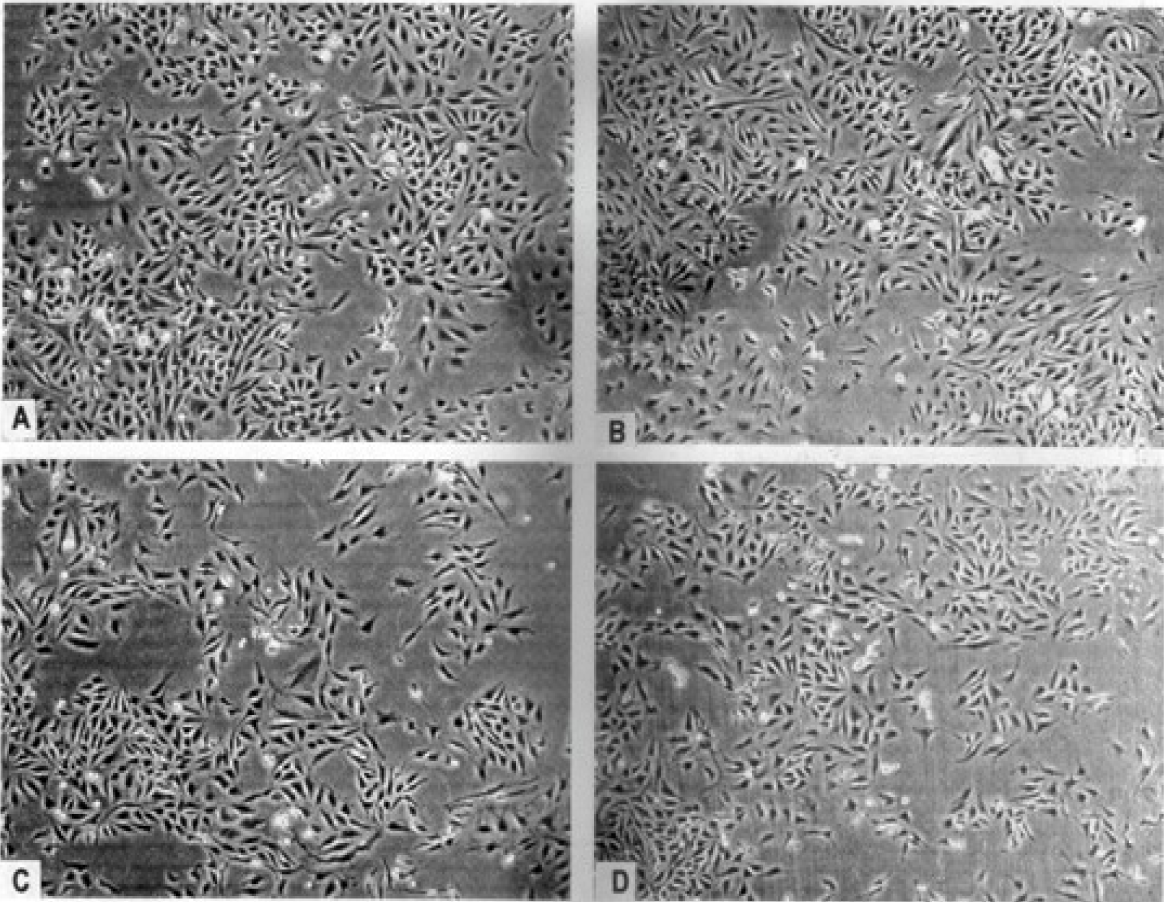
Fig.4B. Toxicological assay of NU2 on HeLa human cancer cells. HeLa cells were grown in presence of varying concentration of NU2 (0, 10, 20, 50 µg/ml) and was observed under light microscope at 400X (Panels A-D). No significant changes of confluency were observed up to 50µg/ml NU2. The 1:8 sub-division of trypsinized cells as in (D) were grown again suggesting minimal effects on HeLa cell metabolism and viability.

**Fig.4B.**
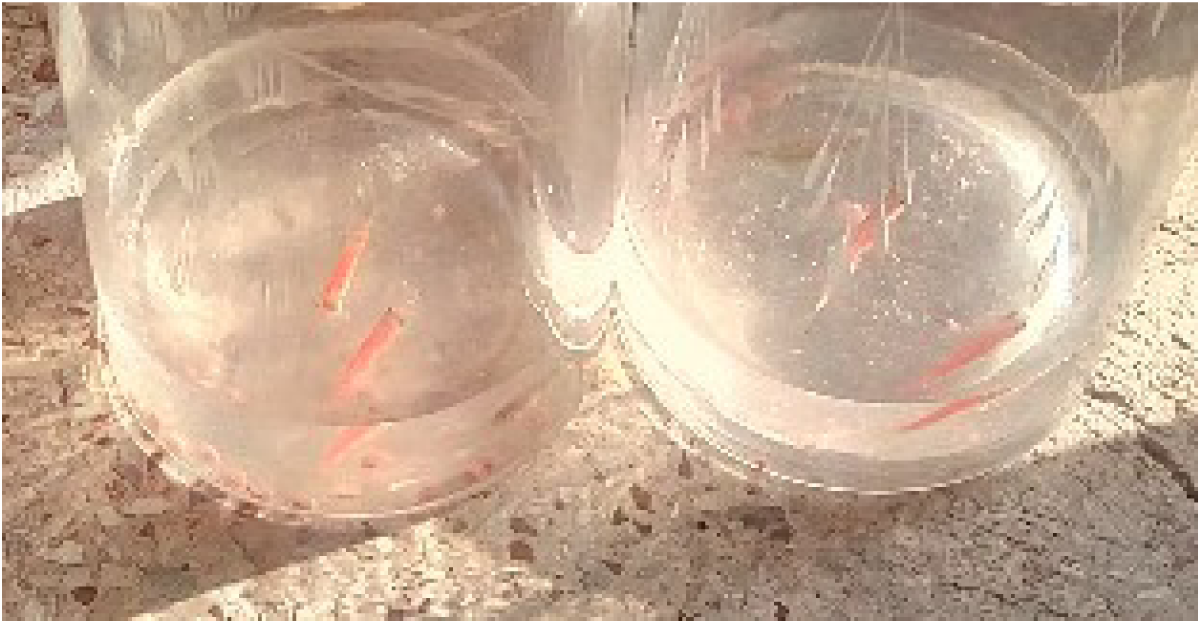
Toxicological study on red Moly fishes. Moly fishes were kept on Kolkata municipality water and 20ug/ml NU2 was added and grown for 3 days. No inhibition was found at that NU2 concentration.

### Elementary analysis of NU2

Elementary analysis was done in a Perkin Elmer CHN Analyzer Model 2400 series II at Indian Association for the Cultivation of Science. The data was not impressive as Carbon= 6.34, H=3.35 and N=0.03 %. There is no nitrogen that was understandable and the NU2 was not an amino sugar. But very low carbon means the compound is very much high oxygen content and likely a poly-phosphate or halogen derivative. The halogen derivative was not possible because that could not bind to oxygen to be necessary for higher percentage of O_2_. The MASS spectra data then told the rest to obtained a 98mu deviant bands for phosphate ions.

### Biochemical assays of NU2

Biochemical assays predicted that NU2 was a glycoside (figure-5). We did find flavones in NU2. As later we found the NU2 compound likely a cyclic polyphosphate glycoside, the biochemical assay of triterpene might be assumed as false. Thus, we predicted a cyclic nature of NU2 to give a false terpenetine assay.

**Fig.5.**
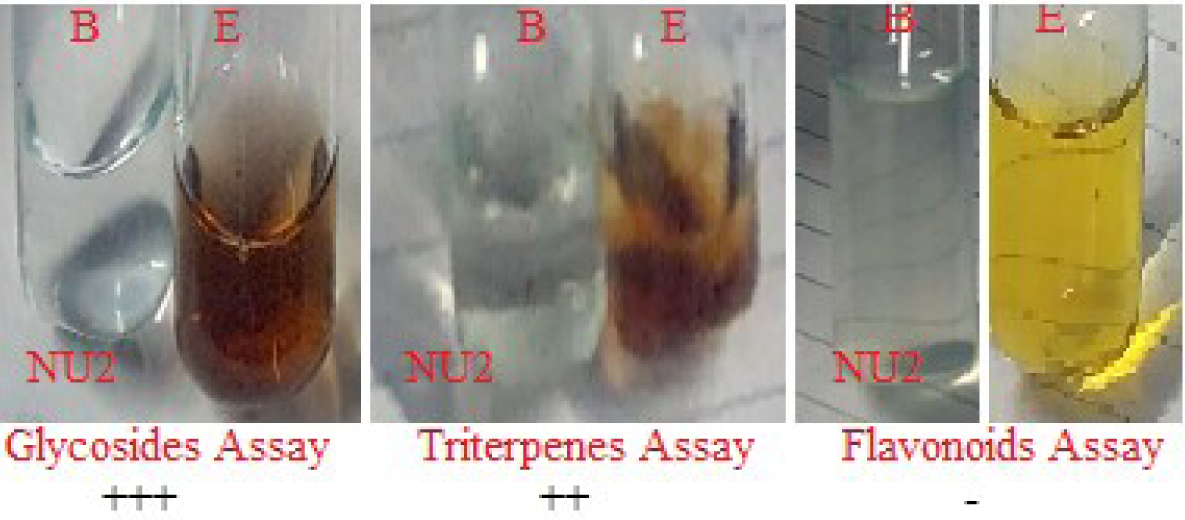
Biochemical assay of phytochemical NU2. It was very positive for glycosides and triterpenes but negative for flavones. It was shown that NU2 gave tittle false test for triterpene possibly due to cyclic structure of NU2 fluoropolyphasphate-glycoside.

### Determination of Molecular Weight of NU2

NU2 was shown to be 454mu mass with formula C_4_O_16_P_4_FH_7_ and next high molecular sub-fragments were 356mu, 297mu and 19mu as judged by MASS spectra (figure-6B). The presence of Fluorine isotopes (F19 and F18) was detected in duplicate bands at most fragments because Phosphorous isotopes (P32 and P31) were very short lives and O16 was abundant only in nature. The NU2 glycoside should contained four phosphate groups and likely one Fluorine atom. The phosphate deviation bands were confirmed by subtracting 454.23mu with 356mu to give 98mu which is molecular weight of phosphoric acid [OP(OH)3]. Thus, we proposed NU2 structure as a Cyclic Fluoro-PolyPhosphate glycosider. NU2 is a wonderful molecule to have a bacterial DNA topoisomerase I target and our finding was novel because there was no good inhibitor to study bacterial DNA topoisomerase IA. Ideally, plant alkaloid and anticancer drug camptothecin was extensively studied by docking computer simulation. Previously, we solved the structure of CU1 isolated from bark of another medicinal plant called golden shower or *Cassia fistula*, which was a poly-bromo derivative of polyphenol likely linked to a tri-terpene derivative. Interestingly, we found CU1 as RNA polymerase target and also inhibited RNA polymerase of *Mycobacterium tuberculosis.* Later, we will show that the NU2 potentially inhibits the relaxation activity of *E. coli* DNA topoisomerase I.

**Fig.6A.**
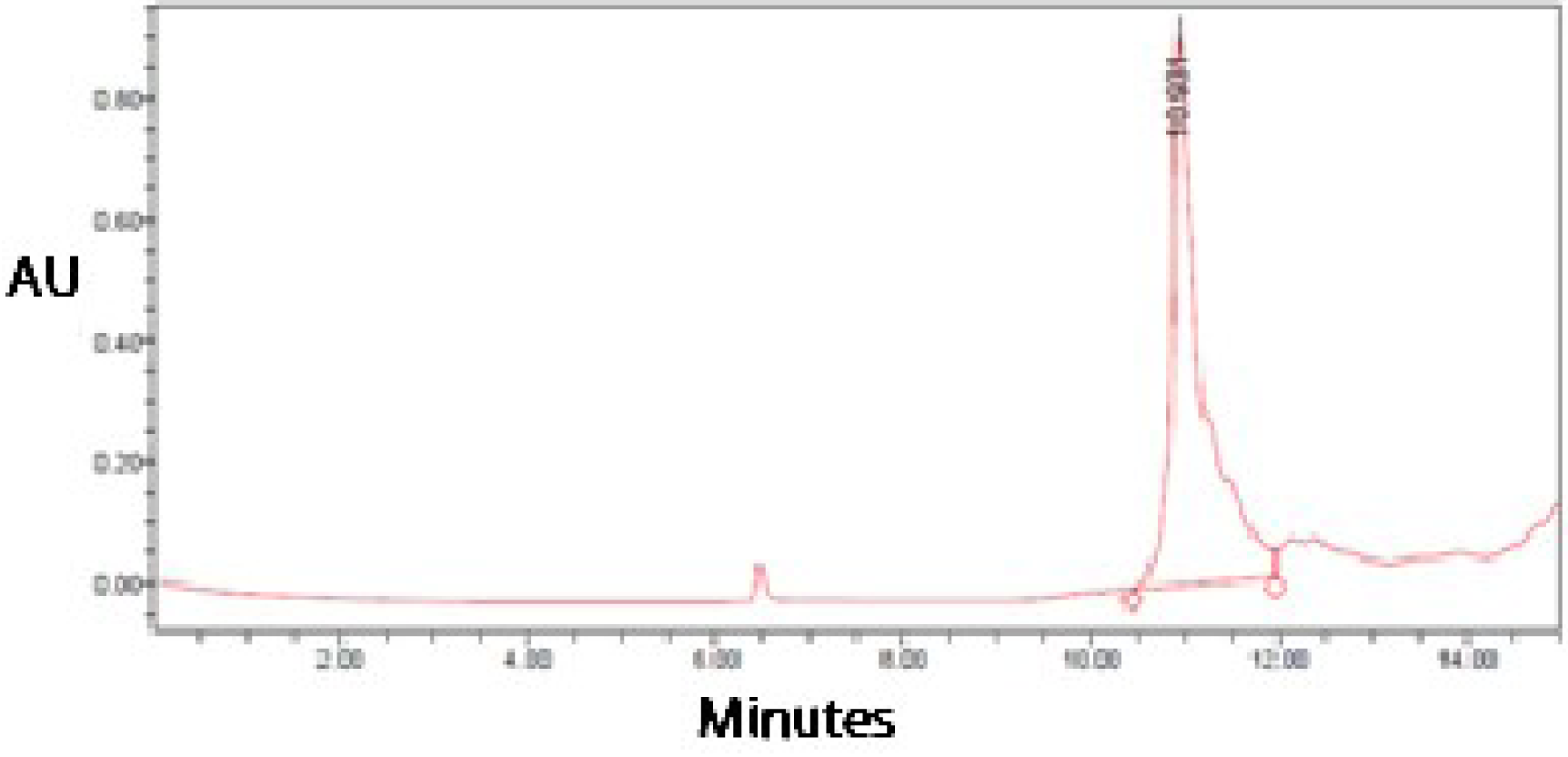
Elution profile of NU2 in HPLC c18 column. The NU2 was eluted at 10.9min whereas we previously showed that CU1 eluted very fast at 3.5min.

**Fig.6B.**
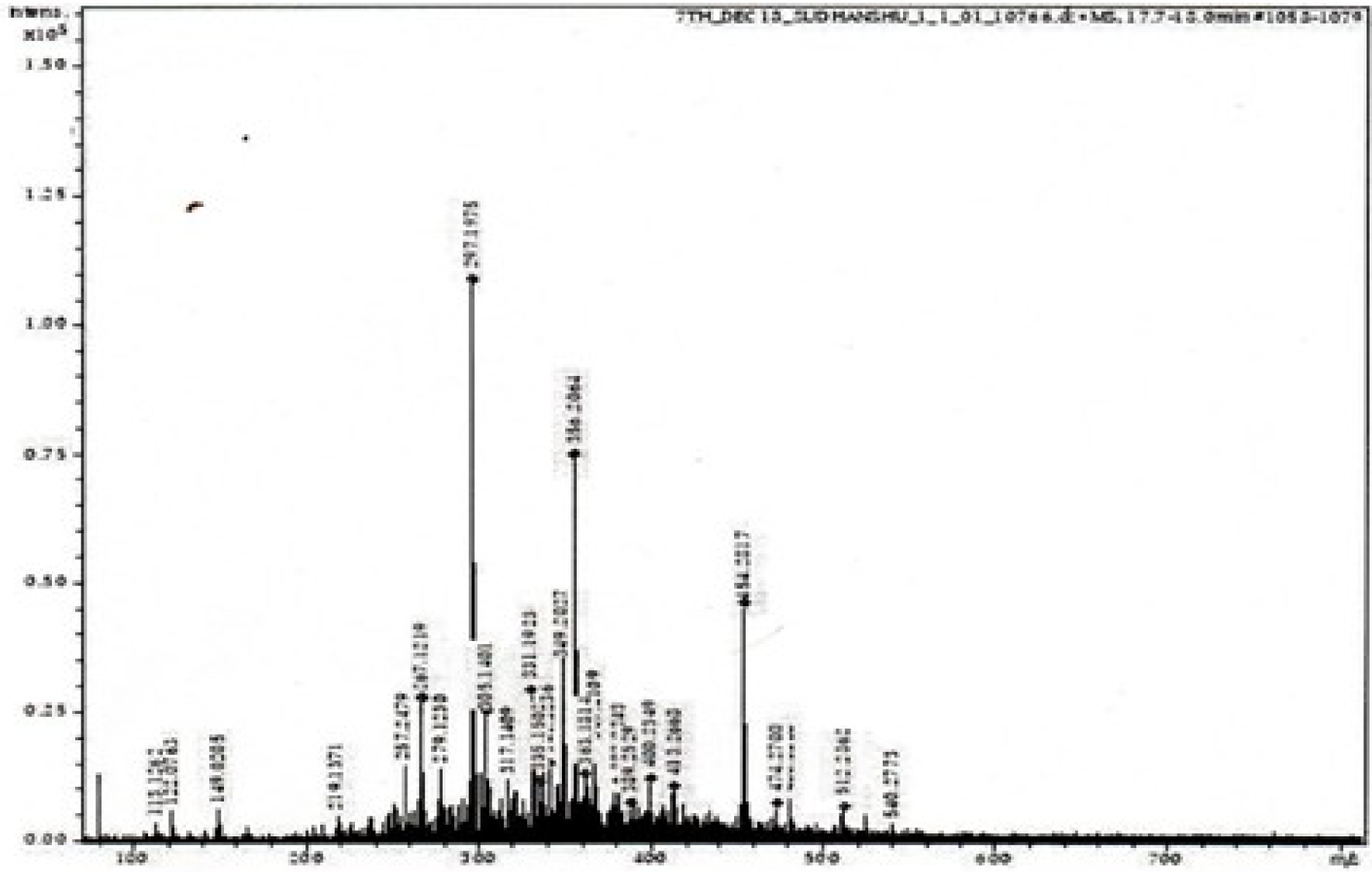
MASS spectra of NU2. It showed NU2 mass as 454.32mu whereas 356.2mu, 297.19mu and 19mu (Florine) are important sub-fragments [Das et al., 2022]. This MASS was done with one time TLC purified and contained some impurities.

### FT-IR spectroscopies of NU2

FT-IR spectra (figure-7A) was highly predicted presence of many -OH group (C-OH and P-OH) as judged by broad range absorption at 2900-3650 cm^-1^ (-OH stretching) with peak at 3443.6 cm^-1^ and C-H symmetric stretching at 2853.8cm^-1^. Secondary alcoholic >C-O-H bending and C-O-stretching might be seen at 1559cm^-1^ and a peak at 1640cm^-1^ likely for water molecules. The 1800-1500cm^-1^ region known as double bond stretching (C=O) but C-H bending could be near 1559cm^-1^. The distinct phosphate group absorption was confirmed at 1021.4cm^-1^. Similarly, skeleton minor vibration at 525cm^-1^ for phosphate group and C-**O**-**P** stretching at 928.7cm^-1^ were determined. The P=O stretching was reported near 1250cm-1 but we found very weak band at that position and found medium band at 1338cm^-1^. However, the P-O-absorption of phosphate ester was aligned at 805.6cm^-1^ for P-O-O-P. The absorption for C-F was reported at 1400-1000cm-1.

**Fig.7A.**
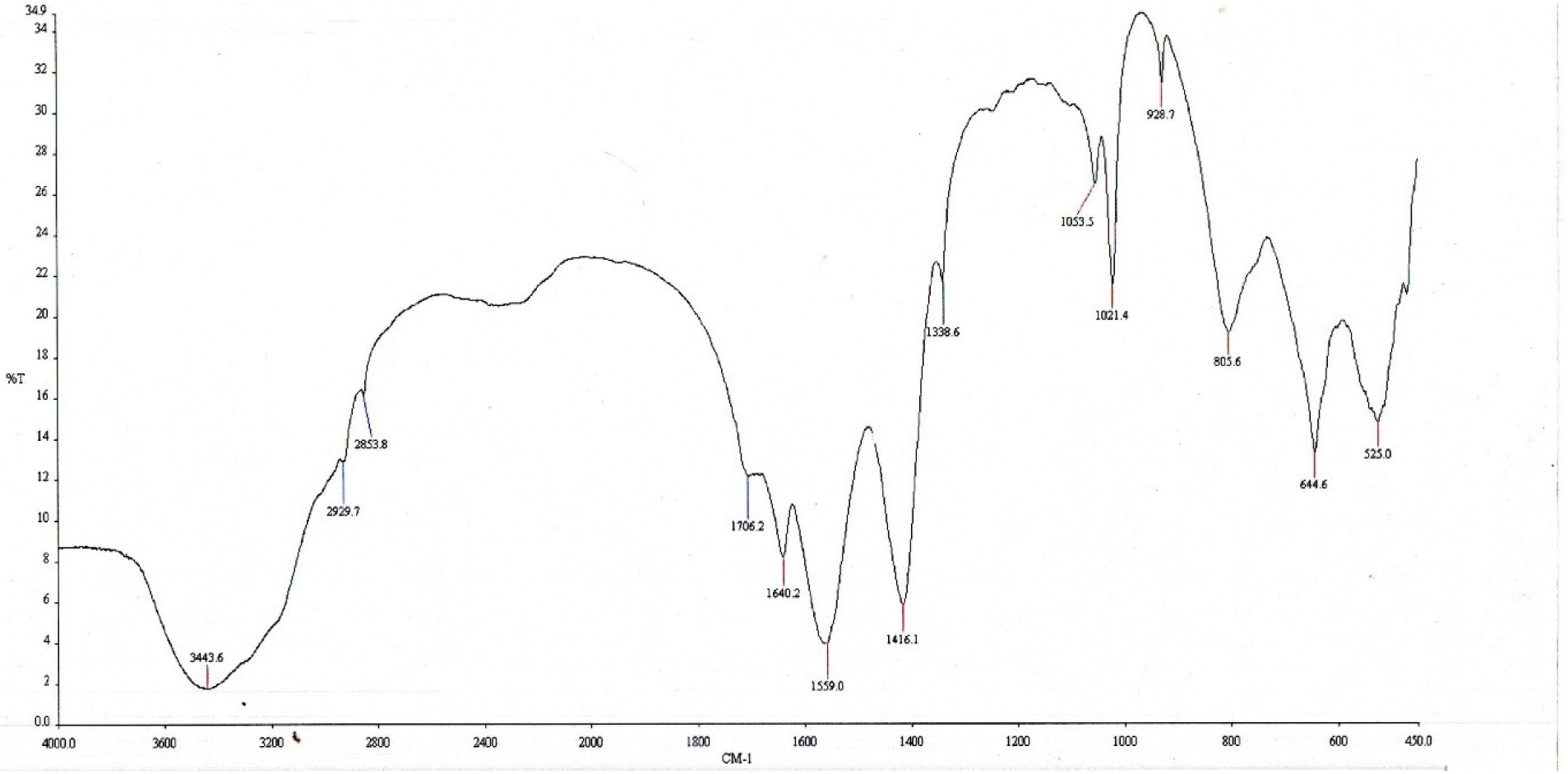
FT-IR spectra of NU2. Many absorption-spectral lines for functional groups obtained: at 2900-3600cm^-1^ for -OH group stretching, 2853cm-1 for C-H stretching, at 1021cm-1 for phosphate stretching, 1559cm^-1^ -C-H bending, at 1416cm-1 for ring -OH, at 928cm-1 for C-O-P ether bond and at 525cm-1 complex structure of polyphosphate were important.

### NMR-Spectroscopy of NU2

Proton-NMR spectra (figure-7B) selected huge absorption at δ=2.269 for -O-H, δ=1.546 for - CH-O-P and -CH2-O-hydrogens and δ=1.292ppm for -CH_2_-. A weak absorption at δ=5.32ppm may reflect primary alcoholic -O-H in glycoside whereas at δ=7.26ppm surely a CDCl_3_ solvent absorption peak. While Carbon-NMR (figure-7C) only gave good absorption spectra at 30.1ppm for C-H and at 207.01ppm for -HC=O and -C-O-P whereas at 77ppm for CDCl_3_ solvent only. We should not expect more peaks in carbon-NMR due to very low carbon content.

**Fig.7B.**
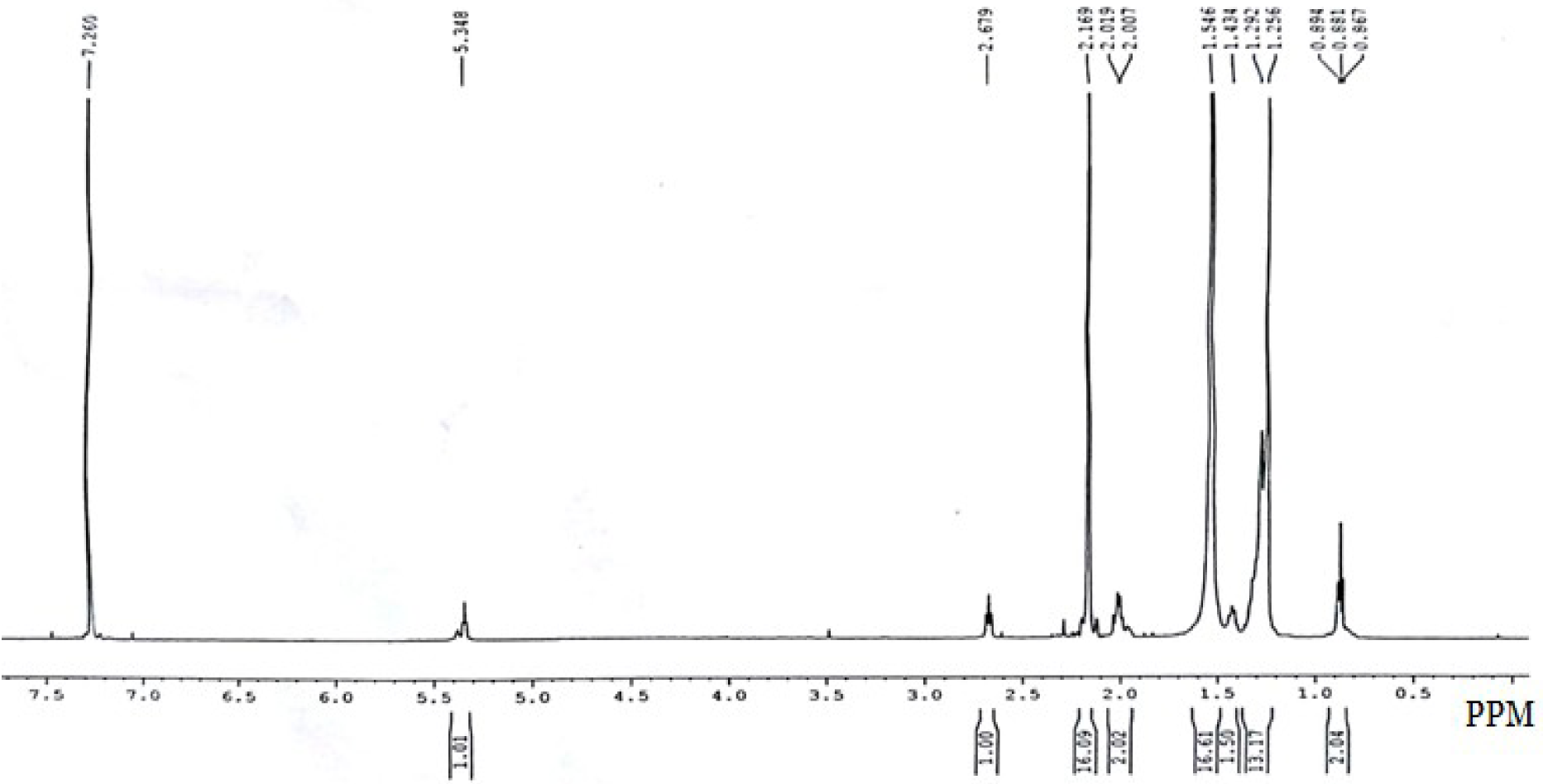
Proton-NMR spectra of NU2. Proton-NMR produced absorption at δ=2.269 for -O-H, δ=1.546 for -CH-O-P and -CH2-O- and δ=1.292ppm for -CH_2_-.and at 7.2ppm for solvent CDCl_3._

**Fig. 7C.**
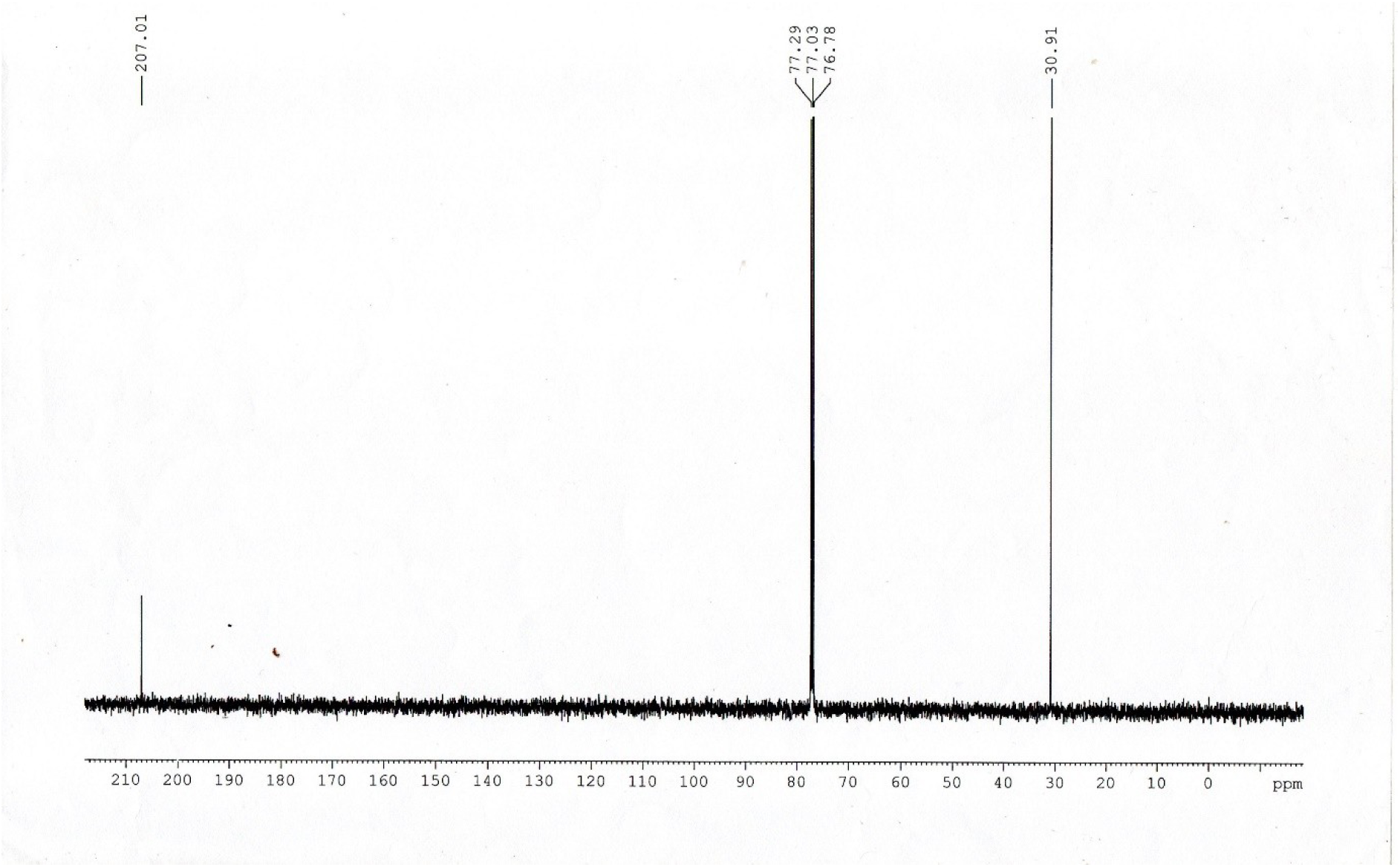
Carbon-NMR spectra of NU2. Carbon-NMR produced 30.91ppm for C-H and 207.01ppm for -CH=0 and -C-O-H absorption spectra. Band at 77.03ppm is for CDCl_3_ solvent.

### Molecular target determination of NU2

DNA topoisomerase I was assayed by reduced mobility of pBR322 plasmid relaxed topoisomers on 1% submerged agarose gel at reduced voltage 15-20V using 20 cm long agarose gel for 8-12 hrs. We found such inhibition of *E. coli* DNA topoisomerase I with 10-20 µg/ml NU2 (figure-8A) whereas JU1 and JU2 phytochemicals from *Jatropha gossipifolia* did not inhibit DNA topoisomerase I (figure-8B). Neither, CU1 from *Cassia fistula* bark ethanol extract has any inhibitory activity on *E. coli* DNA topoisomerase I. Abundant JU2 phytochemical from *Jatropha gossipifolia* roots did not show any RNA polymerase activity either (in preparation). We have not tested DNA gyrase, DNA topoisomerase III and human DNA topoisomerases (I, II and IV) due lack of resources but such study should be done for future for drug development strategy against human. We had previously shown that CU1 phytochemical from *Cassia fistula* has strong RNA polymerase target but CU3 phytochemical from same plant showed no RNA polymerase inhibition. Moreover, No DNA polymerase activity was demonstrated both for CU1 and NU2. We have also tested NU2 on RNA polymerase. NU2 was a very weak *E. coli* RNA polymerase inhibitor but no such activity was found in *M. tuberculosis* RNA polymerase and thus we have not followed such study further (data not shown).

**Fig.8A.**
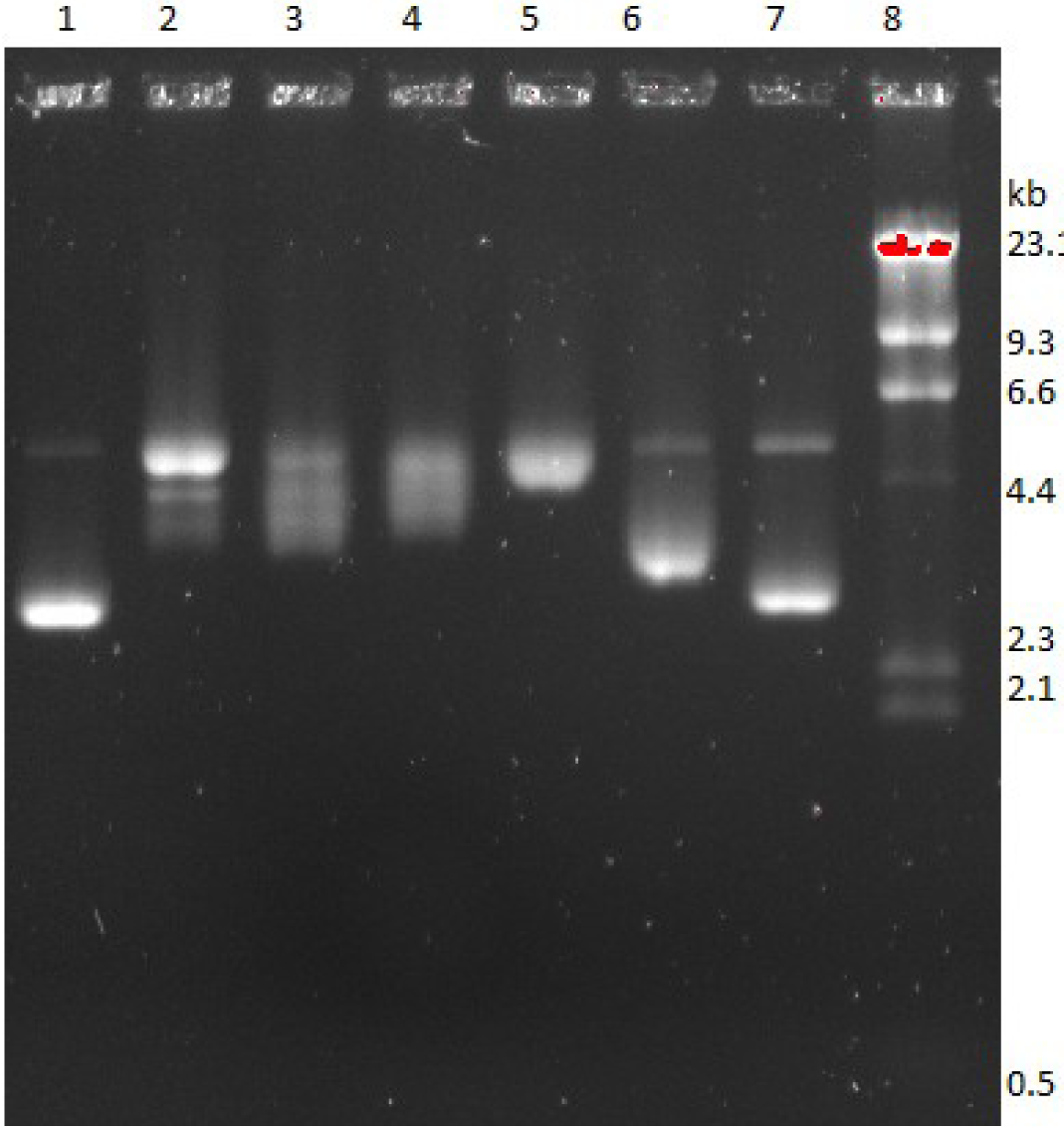
*E. coli*DNA topoisomerase I inhibition by NU2 phytochemical of *Suregada multiflora*. Lane 1, control 0.5µg supercoiled pBR322 plasmid DNA, Complete assay with 1unit topoisomerase I, lanes 3, 4, 5: +0.4, 0.6, 0.8unit enzyme., lane 6, +10µg/ml NU2 and lane 7, +20µg/ml NU2. Lane 8, Lamda HindIII DNA marker. It appeared NU2 very strong bacterial DNA topoisomerase I inhibitor.

**Fig.8B.**
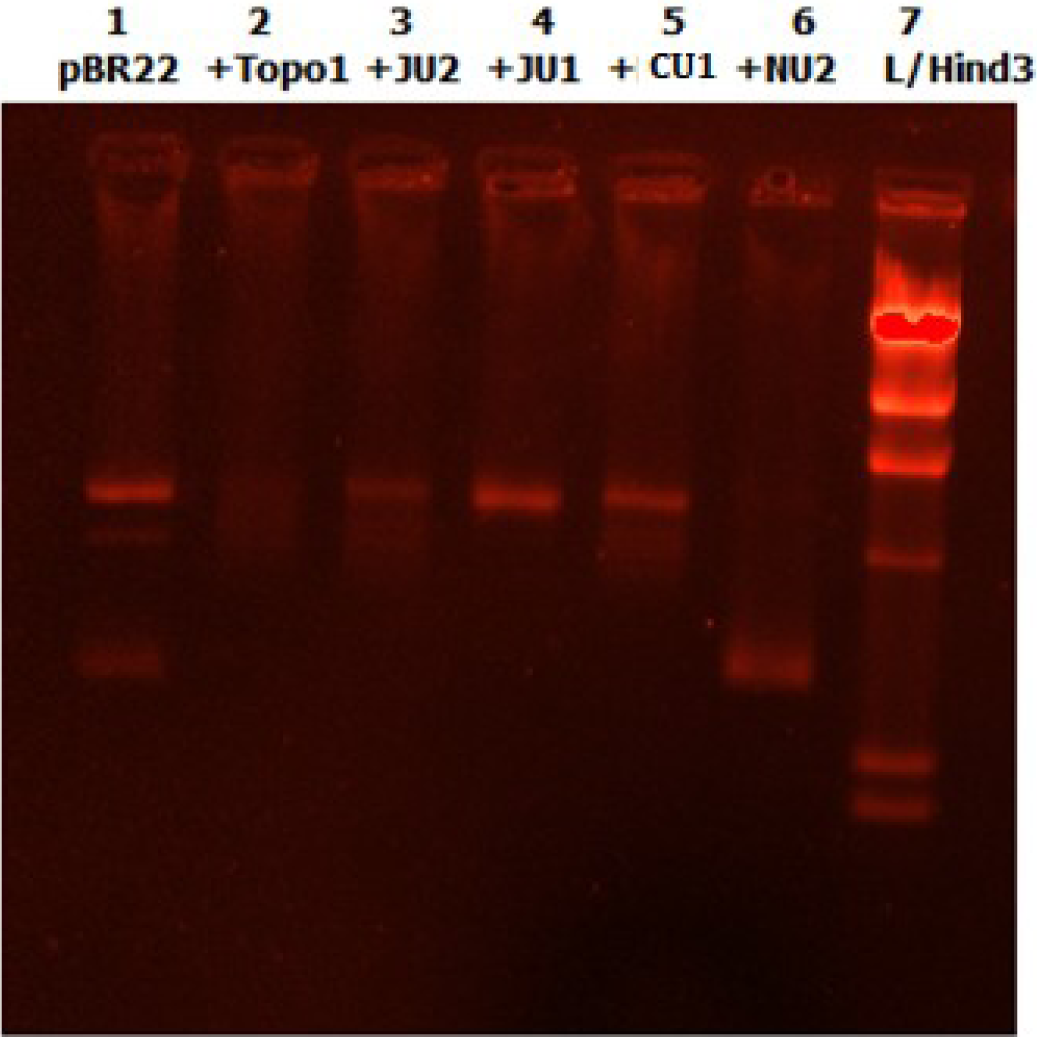
Compare of *E. coli* DNA topoisomerase I inhibition by *Suregada multiflora* and *Jatropha gossipifolia* different phyto-chemicals. Lane 1, 0.2µg pBR322., Lane 2, +0.5unit enzyme, lanes 3, +JU2, lane 4 +JU1., Lanes 5, +CU1, Lane 6, +NU2 each 20µg/ml. Lane 7, Lamda HindIII DNA marker. DNA topoisomerase I specifically inhibited by NU2.

### SwissDock Computer simulation to study DNA topoisomerase-drug interactions

Next, we studied the molecular drug-enzyme interactions using SWISSDOCK. The structures NU2.1, NU2.2 and NU2.3 (figure-9) were drawn using Stacher of SwissDock to give distinctive SMILES. The *E. coli* DNA topoisomerase I (PDB ID: 1MW8) modelling was shown in figure-10 where NU2.3 bound to different domains dominated with basic amino acids. The NU2 binding was very specific at the domain-1 of *E. coli* and *L. donovani* topoIII (figure-11A). We have done kinetics of Mycobacterial topo1A with NU2 and the drug can bind at different domains with lower efficiencies (figure-11). However, such interactions with low efficiency were also found with eukaryotic type-2 DNA topoisomerases (data not shown). Although, we found that human topo3b had some specificity to NU2 (figure-12A) while unicellular human macrophage parasite *Leishmania donovani* topo3 bound at the domain-2 very similar to bacterial topo3 (figure-12B). TB is a central problem in India. Although NU2 bound to topo1 of *M. tuberculosis* (figure-13) but surface amino acids could not be aligned to primary sequence (protein id. 8TFG_A). Truly, more topoisomerase crystal structures to be created to get a conclusive evidence that NU2 specifically binds to topo3.

**Fig.9.**
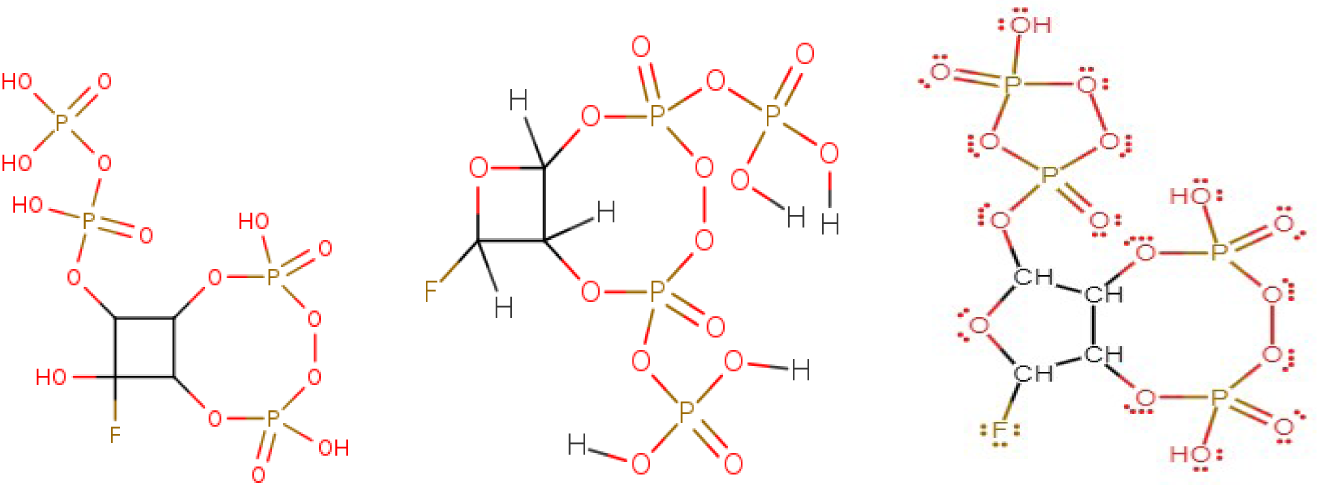
Proposed structures of cyclic glycoside polyphosphates (NU2) that used for SwisssDock against DNA topoisomerase I and DNA polymerase III. The structure was made assuming oxaloacetate or glyceraldehyde 3-phosphate condensed with phosphate to make NU2. The molecular weight of NU2 was 454 kDa with low carbon content (16%) but high phosphate content and one Fluorine atom were considered.

**Fig.10A.**
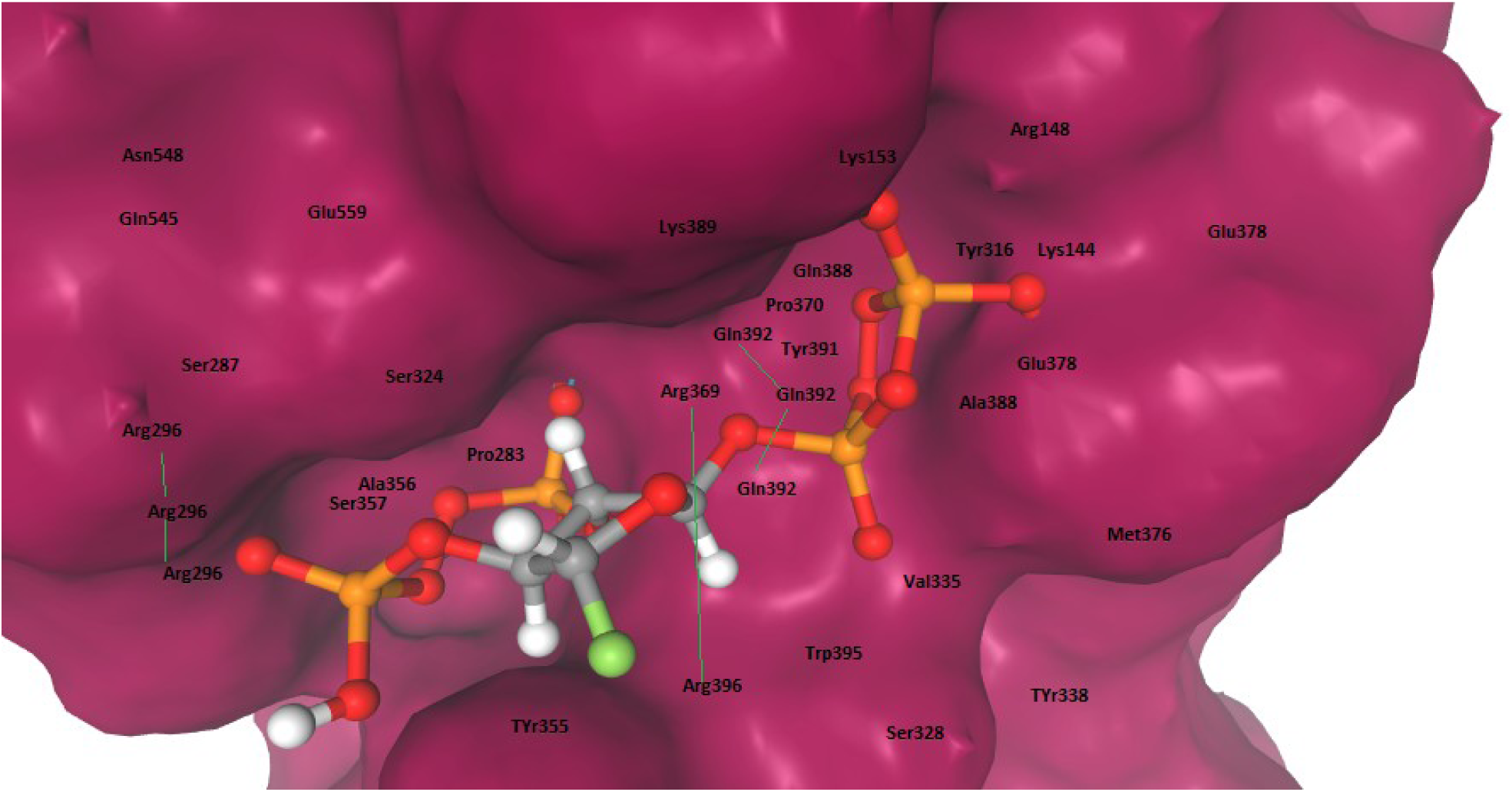
Binding of NU2.3 at the domain-2 of *E. coli* DNA topoisomerase IA (PDB ID: 1MW8). The Lys144, Arg148, Arg296, Tyr316, Ala356, Tyr355, Ser357, Arg369, Gln378, Arg396 may be important amino acids to build the NU2.3 binding domain. However, Arg144, Thr316, Tyr355, Arg369 and Tyr391 are conserved in *M. tuberculosis* DNA topoisomerase IA. The positions based on BLASP-2 between sequences 1MW8 (*E. coli*) and 8TFG (*M. tuberculosis*) DNA topoisomerase IA amino acid sequences.

**Fig.10B.**
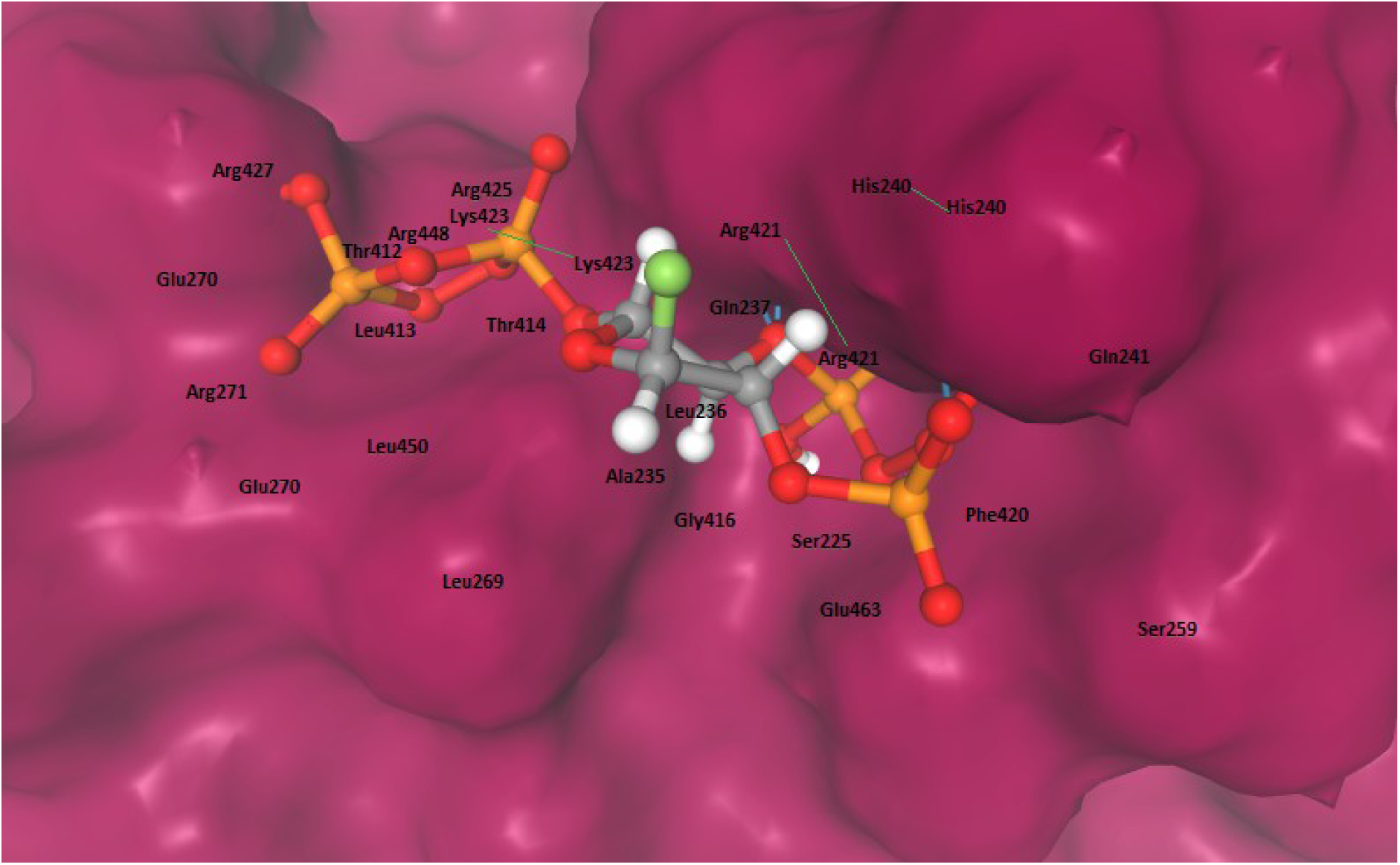
Binding of NU2.3 at the neighbouring of active domain-2 of *E. coli* DNA topoisomerase I (PDB ID:1MW8). The Ser225, Leu236, Gly237, His240, Gln241, Glu270, Arg271, Arg421, Arg425, Arg427, Leu413, Thr414, Gly416, Phe420, Lys423, Arg448 may be involved in building active domain to bind NU2.3. However, Ser225, Leu236, Gly416, Arg427, Arg448 are conserved in *M. tuberculosis* DNA topoisomerase IA.

**Fig.10C.**
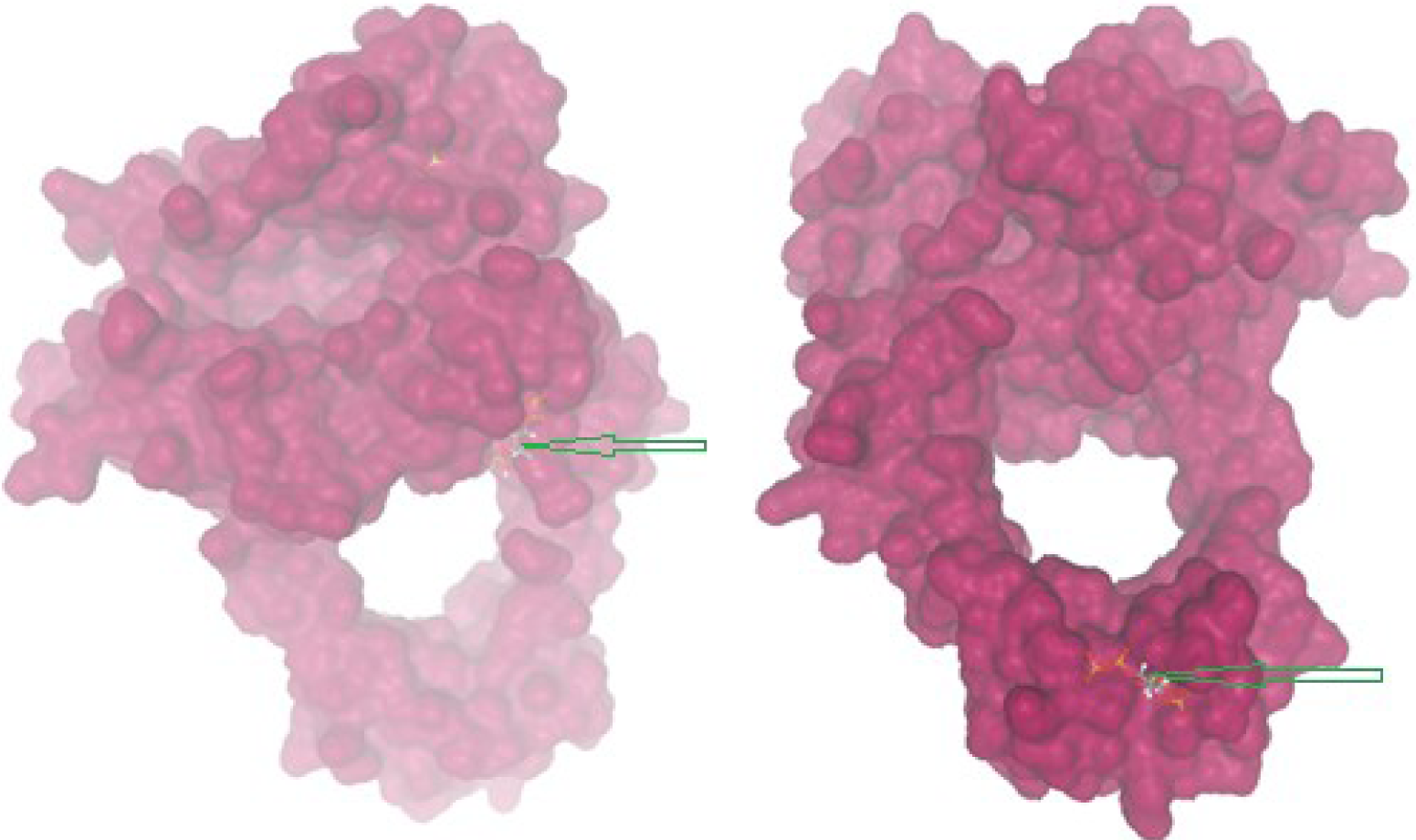
Localization of multiple binding sites of NU2.3 at the 3-D surface structures of *E. coli* DNA topoisomerase IA (PDB ID: 1MW8) binding to NU2.3. During simulation at the active site (Box centre=28/38/30) active-binding site located at the surface as in figure-10A (left picture, see middle arrow) while during simulation at the domain 4 (Box centre=33/24/-15) active binding site (right picture) located at the other site (see down arrow). This predicted multiple binding sites on *E. coli* DNA topoisomerase IA.

**Fig.11A.**
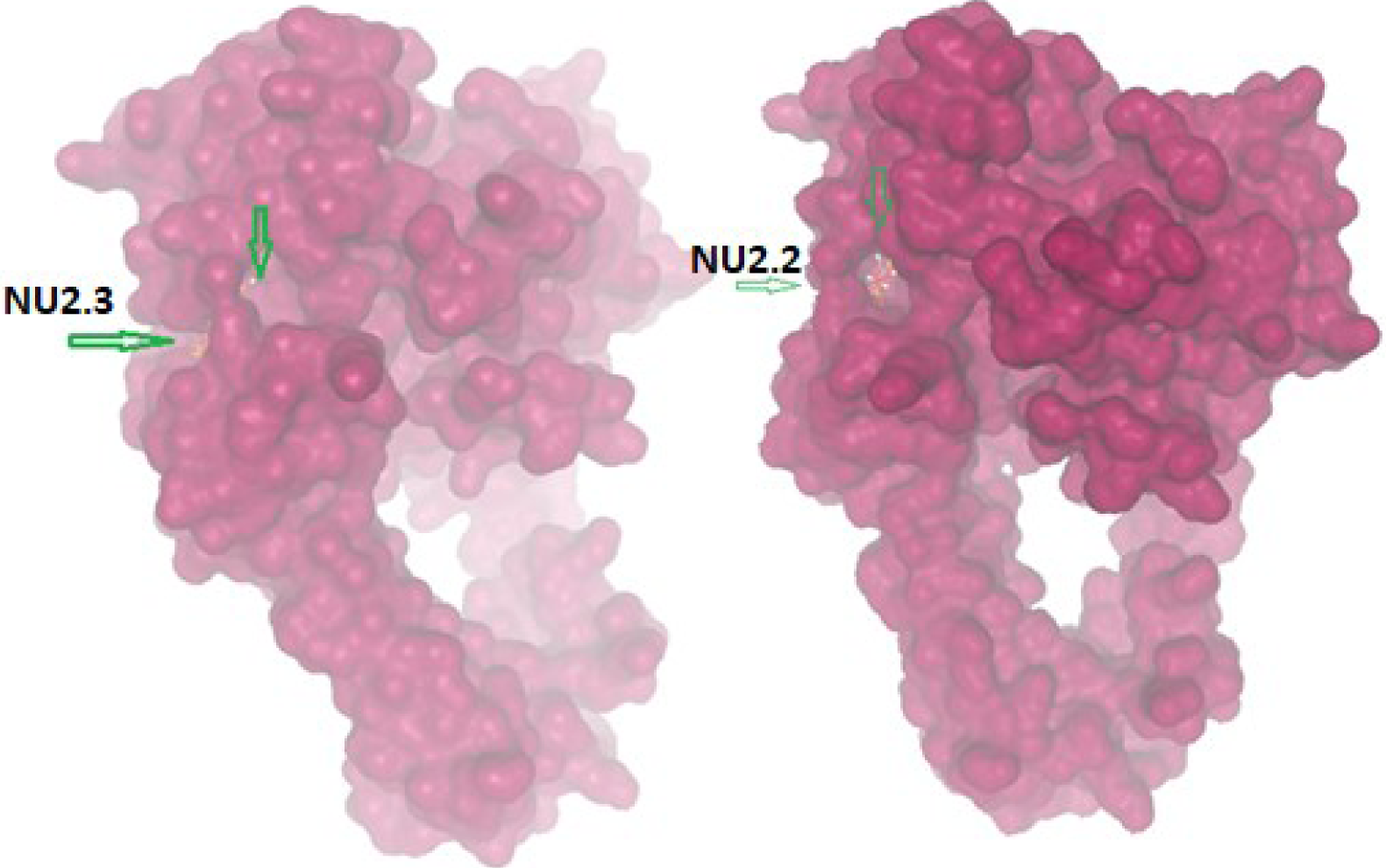
Binding of NU2.3 and NU2.2 to *Escherichia coli* topoisomerase III (PDB ID:1D6M). Both drugs bind to the same active site. The AC score= -236.69 and SwissParam score= -7.73 for NU2.2 while AC score= -238.83 and SwissParam score = -7.91 for NU2.3 targeting *E. coli* topoisomerase III. This binding occurred in the groove of active site and suggested NU2 drug as strong inhibitor of topoIII than topoI, being both type-I DNA topoisomerases of *E. coli*.

**Fig,11B.**
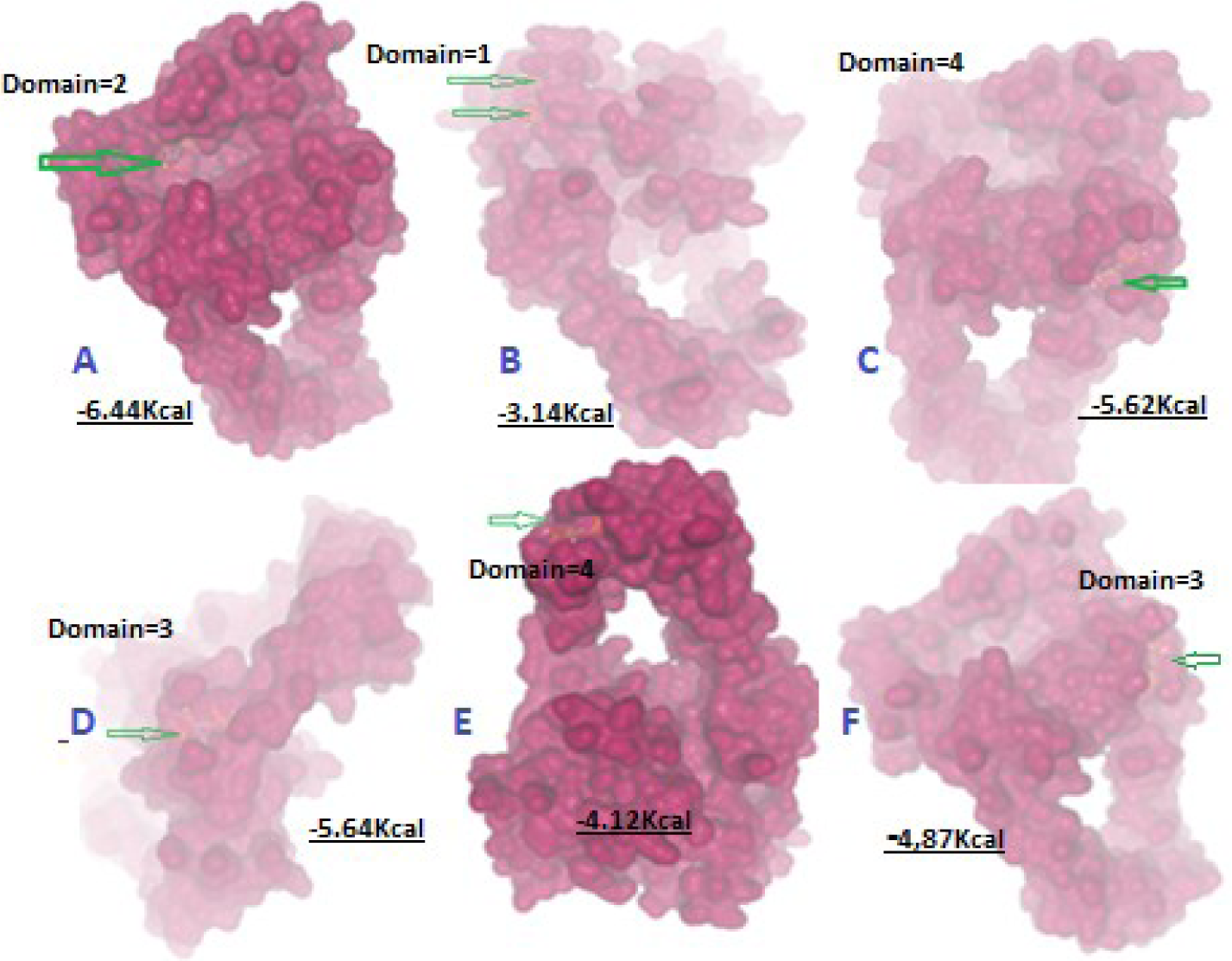
Kinetics of Binding of NU2.3 fluoropolyphosphate-glycoside. [SMILES= OP1(=O)OOP(=O)(OC2OC(F)C3OP(O)(=O)OOP(O)(=O)OC23)O1] to the different domains of *Escherichia coli* topoisomerase III (PDB: 1D6M) using software AutoDock-VINA. The above binding energy of -6.0 Kcal/mole was taken as potentially important drug-enzyme interactions that might be inhibited the DNA binding, nicking, swivelase and ligation reactions.

**Fig.12.**
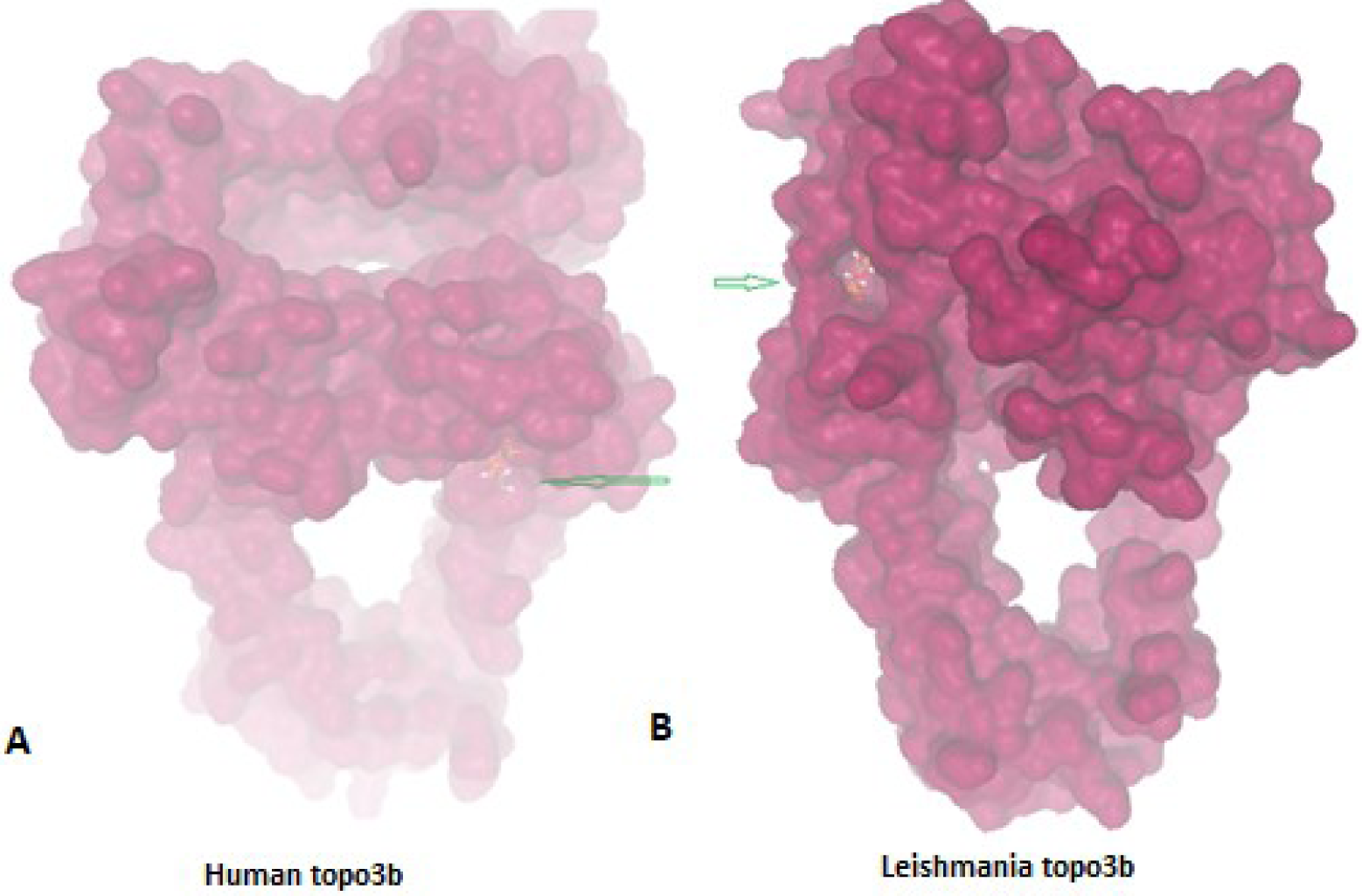
**(A) Binding of NU2.2 to the human topoisomerase 3b1 (PDB ID:9C9Y).** The AC score= -410.93 and SwissParam=7.35. It is not similar to binding of NU2.3 to *E. coli* DNA topoisomerase I (PDB ID:1MW8). **(B) Binding of NU2.3 to *Leishmania* topoisomerase 3b (PDB ID:1D6M).** The AC score (CHARMM force field energy and solvation energy) = -235.69 Kcal/mol and SwissParam score (free binding energy) = -7.73 Kcal/mol. The free binding energy in the range above -8.0 was considered as good drug-protein interaction.

**Fig.13.**
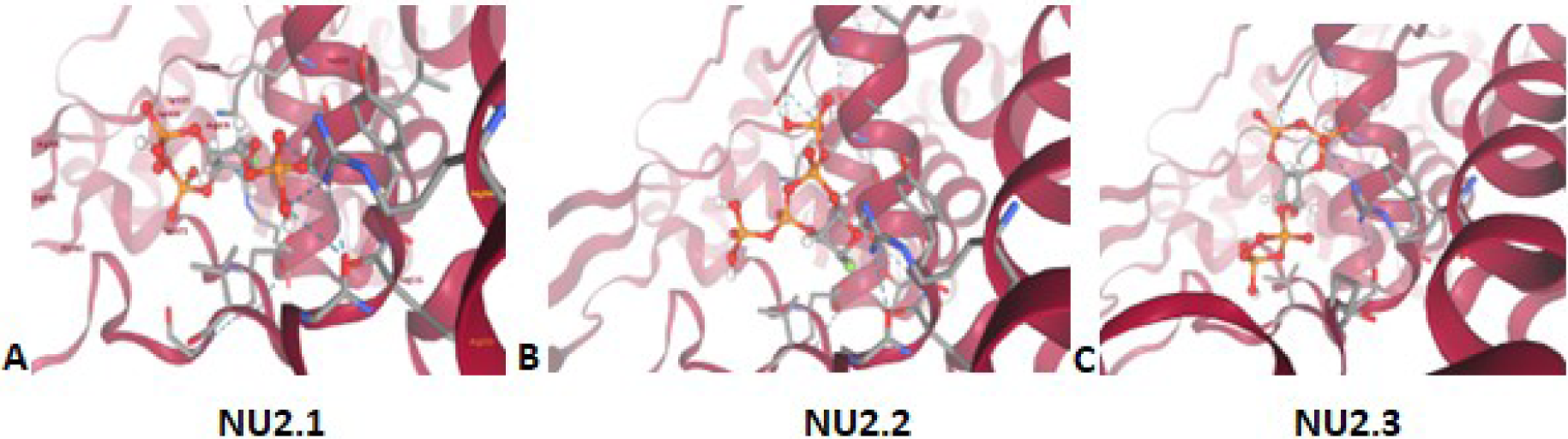
Binding of NU2.1, NU2.2 and NU2.3 3 at the domain-2 of *M. tuberculosis* DNA topoisomerase I (PDB ID: 8TFG). Active binding surface amino acids are: (A) Lys185, Arg318, Thr474, Asp476, Glu477, Val479, Gly480, Ala515, Tyr517, Thr518, Arg516, Ser521; (B) Arg206, Ala205, Trp561, Phe564, Ala565, Asn662, Ile664 and (C) Trp561, Val562, Ala565, Arg658, Asp686, Lys661, Asn662, Tyr665. But we could not able to match the primary amino acid sequence of *M tuberculosis* topoI which is very easy for *E. coli* topo1 and topo3.

## Discussion

Antibiotic monopoly to treat microbial infections was started in 1940s after World War II. But during 1960s onwards huge increase of multidrug-resistant infections suggested the end of antibiotic age [Bassetti, 2009]. Most early antibiotics are useless in MDR infections singly or in combination and in a single plasmid as high as 15 *mdr* genes were reported (Lei, et al., 2017]. Still, study suggested that combination old antibiotic might be good in MDR skin infections. Major reasons of AMR are the accumulation of early *mdr* genes (blaTEM1, blaOXA1/2, *tetB, cat, sul1/2, strA/B*) in small R-plasmids [Projan and Novick, 1988; Chakraborty, 2018]. But use of higher derivatives of antibiotics also created conjugative plasmids with new derivatives of *mdr* genes *(blaOXA51, blaCTX-M-1/2/9, blaNDM1, aphA4, aadA2, mcr-1* [Huang et al., 2013]. Further, target mutations (rRNA, porins, PBPs, gyrA/B) of chromosomal genes and activation of multi-drug efflux proteins (tetA/C, acrA/B, mexA/B, macAB, mtrCD) acted synergistically today to void many antibiotics actions [Delmar., 2014; Lynch, et al., 2013]. We checked the huge MDR situation in Kolkata Ganga River superbugs as well as chicken meat and milk. We clearly showed studying plant-derived heterogeneous phyto-antibiotics as described in ancient Hindu Civilization and in Sanskrit books like Charaka Samhita, Satruta Samhita and Atharva Veda, might be good to cure todays MDR infections due to development of newer methods to purify and to characterize the structure and function [Chakraborty, 2015; 2017a,b; Chakraborty et al, 2022].

We clearly demonstrated in this article that *Suregada multiflora* glycoside-polyphosphate actively an inhibitor of DNA topoisomerase IA. However, we unable to check any activity on DNA gyrase or eukaryotic DNA topoisomerases due to funding problem and my retirement. Further, COVID-19 situation greatly impacted the project. Interestingly, a wide range of MDR-bacteria isolated from Ganga River water, chicken meat, milk, and human hair were inhibited by the four times TLC-purified NU2 as well as concentrated ethanol extracts of *Suregada multiflora* bark and root. Further, we showed that cultivation in Kolkata flat gave highly effective phytochemical to inhibit the MDR bacteria and root culture further increased the bioactivity. Thus, the report is very important for drug discovery scientists. Moreover, NU2 is abundant and any drug company can easily formulate the ayurvedic drug in crude extract or in TLC purified form which should be >90% pure. Important phyto-drugs like artiminisin, taxol, reserpine, quinine, camptothecin were used today with good promise to treat cancer, malaria, sleeping sickness, and bacterial sepsis.

MDR-bacteria pollution is havoc in water similar to other chemical pollution worldwide [Chakraborty & Roy., 2018]. Ganga River of India is highly polluted and other famous rivers like Cuyahoga River (Ohio, USA, oil release), Potomac River (Maryland, USA, Industry sewage); Pripyat river (Japan, Chernobyl disaster in 1986), and Songhua river (China, 100 tons toxin release in 2005) have also listed as very dangerous to bath and consume fish from it. Truly, deadly bacteria could be generated in such polluted river. Accumulation of metal drug efflux genes in MDR bacterial plasmids was a reason for such consequence (Chakraborty, 2020). If we estimate average 120gm of feces per day, and 1/3 of Indian live in the cities located in the Ganga River bank, then 40,000,000 KG of feces are released into Ganga River with ∼5×10^14^ uropathogenic and MDR bacteria per day [Chakraborty., 2015., 2017b]. In other estimate, a 160 pounds person produces a pound poop giving 127KG poop/person/year and 900 million tonnes poop produced worldwide. An average people release 1200ml urine with average bacteria 2×10^3^ cfu/ml. So, Ganga River may receive 40×10^7^ x 2×10^3^ x 1200x 365=3.5 x 10^17^ bacteria/year. Thus, toxicity and multi-resistance appeared ubiquitous [Fosberg, et al., 2014; Chakraborty., 2017c.,2017d]. We got so many dangerous pathogens in Ganga River water that we stopped the project in fear of unknown infections.

Importantly, 2×10^12^ intestinal bacteria in the intestine are constantly synthesizing 20 vitamins and complex bio-molecules for our body and one high dose of antibiotics is enough to kill all such microbiota [Dethlefsen et al., 2008]. Thus, it appears, multidrug-resistant genes creation is to protect microbiota from repeated doses of antibiotics that we have consumed since 1940s [Chakraborty, 2020]. Thus, MDR bacteria will be the resident of intestine favouring vitamin biosynthesis, needed for normal human metabolosome. Surely, such symbiosis is so strong that vitamins synthesizing genes are accumulating in plasmids to save human and animal. High quality research from US Human Microbiome Project (HMP), European Metagenomics of the Human Intestinal Tract (MetaHIT) and others have demonstrated the beneficial functions of the normal gut flora (>35000 species) on health [Dethlefsen, et al., 2008]. It is thus G-20 Nations in Germany are united for active research on MDR bacteria to stop superbug horror. Further, secondary MDR bacterial infections in COVID-19 infections are a threat to human race [Tiri, et al., 2020; Vaillancourt, et al., 2020; Chakraborty, 2022]. Interestingly an improved MDR-Cure organic phyto-extracts (*Cassia fistula*, *Suregada multiflora, Jatropha gossipifolia etc.*) including two spices Labanga and Derchini (*Syzygium aromaticum* and *Cinnamomum zeynalicum)* inhibits Kolkata superbugs and gives a hope for new drug development (Chakraborty, 2017a,b). Still, we should take good bacteria (*Lactobacillus* probiotic*s*) after treatment of multiple strong antibiotic doses and capsule with such bacteria available today.

The NU2 phytochemical inhibited MDR bacteria as well as many unicellular parasites. My grandfather used Narenga plant (*Suregada multiflora*) leaves to relieve the fever of Pali-Azar (malaria) being a traditional antimalarial herb. I was sent 25% ethanolic (1:5w/v) extract of leaves to John Hopkins University School of Medicine (USA) and Dr Shapiro was sent me data that 69% inhibition of *Plasmodium falciparum* obtained *in vitro*. Later I had spent to Johns Hopkins for three months to complete malaria work as well as unique bi-subunit *Trypanosoma brucei* DNA topoisomerase IB work [Boldly, et al., 2003]. NU2 spectral analysis suggested a glycoside-polyphosphate but carbon content was very low (figure-6) and we postulated few structures based on low carbon content, absence of nitrogen and MASS, FTIR and NMR spectral analyses (figure-7).

There are some natural compounds for example; curcumin, resveratrol or quercetin can bind to many target molecules implicated in human disease. Some of these targets such as acetylcholinesterase or monoamine oxidases A and B, are unique to animals and have no homologues in plants that produce these natural agents. Phenolic glucoside arbutin, which is used to treat urinary tract infections itself, is ineffective until it is hydrolysed and oxidized to hydroquinone in the human body. Further examples are the sennosides, which are converted into laxative Anthrones by bacteria in the gut. Similarly, conjugated phytoestrogens have to be hydrolysed in the stomach or the gut to exert their oestrogen-like effects. Strictly speaking, these plant molecules are not drugs, but pro-drugs. Alzheimer’s disease is characterized by the death of nerve cells in the cerebral cortex and is the most common subtype of dementia that affected 25 million people worldwide in 2000 and is expected to increase to 114 million by 2050. Natural products like nicotine, caffeine, resveratrol, quercetin and curcumin have anti-AD properties [Park., 2010; D’Onofrio, et al., 2017]. We hope NU2 cyclic polyphosplate will be a future drug against MDR bacteria but more research and clinical studies to be performed. In this context, NU2 was found less toxic to HeLa human cancer cells as well as Molly fishes. Further, it was found ∼10 times less active than imipenem (a carbapenem drug that inhibits cell wall peptidoglycan synthesis similar to other penicillin drugs) *in vitro* against MDR *Escherichia coli* at low concentration (data not shown). However, meropenem and imipenem were very costly and for high toxicity profile needed hospitalization for treatment. NU2 was very soluble in water than CU1 drug from *Cassia fistula* bark and CU1 showed less active in LB medium against MDR-bacteria [Chakraborty, et al., 2022]. Others also have isolated antifungal and antibacterial agents from Indian medicinal plants [Kumar, et al., 2006; Xu, 2014]. Recently, phage therapy comes of age to tackle MDR bacterial infections as well as different gene therapy technologies [Chakraborty, et al., 1993; Dunbar, et al., 2018; Yu, et al., 2016]. WHO also supports the non-toxic green antibiotics as alternate to synthetic drugs [Das et al., 2022].

We found DNA topoisomerase 1A as target for NU2 glycoside-polyphosphate [Lima et al 1994]. The SwissDock modelling found NU2 binding sites at the active sites of *Escherichia coli* DNA top1 and top3 as well as *Mycobacterium tuberculosis* DNA topoisomerase I (PDB ID: 8TFG). Simulation of binding surface amino acids were matched to the sequence of the E. coli enzymes but not matched with mycobacterial enzyme. This conflict is very disappointing and may have few interpretations. First, the amino acid sequence was modified by site-directed mutagenesis but modified sequence was not reported. Secondly, AI-guided software may be interpreted the crossed amino acids as new entity side chains (R). If it is, then cross over amino acids appeared a new amino acid (like a phenomenon of mutation) in 3-D protein structure and totally over ruled the primary amino acid sequence and genetic integrity. Thus, in such situation, it is hard to find a match between simulated amino acid and primary sequence amino acid. Lee et al have shown that TopoIII also involved in the decatenation of catenenes produced during the replication and previously it has been shown that topoIV not gyrase was also necessary for such function (Lee et al, 2019). We believe from computer simulated graphics that topo3 will be a great target for NU2 whereas NU2 also bound to *Leishmania donovani* topo3, a type I DNA topoisomerase was shown to the sole target of urea stibamine and sodium stibogluconate [Chakraborty & Majumder, 1988; Chakraborty et al., 1994]. Such topo3a binds to phenyl Sepharose at 1.5M NaCl while topo1B was eluted. On the other hand, topo2 was separated on hydroxyapatile column eluted at 0.3M phosphate and topo1B plus topo3 eluted at 0.4M phosphate. On phospho-cellulose column topo2 eluted on 0.4M NaCl while topo1b eluted at 0.8M NaCl. Sadly, still there is a controversy on the purity and structure of the antimonial drugs and search did not find any crystal structure with known PDB ID [Frezard et al. 2009]. We did multi-alignment and Swiss Modelling to pinpoint that NU2 might be also the target of kinetoplastid topo3 [Mondragón and DiGate, 1999]. In human topo3a and topo3b had sequence divergent and multiple mitochondrial topoIb other than nuclear topo1B were suggested. In such scenario, the inhibition of animal cells at 20u/ml drug was not found [Chakraborty, 2017]. We are discussing such points because Kala-Azar and malaria drug development is a priority area in India and DNA topoisomerases have shown as potential therapeutic target. Therefore, to study drug-enzyme interactions, you have to purify the very similar enzymes like topo1A, topo1B, MiTopo1B, topo2, Gyrase (bacterial topo2), topo3, topo4, topo5 (Reverse gyrase, heat stable) to check the action of unknown synthetic chemicals and phyto-chemicals.

*E. coli* topoisomerase I (omega protein, protein id. BAA14811) carboxyterminal 280aa was not included in PDB sequence ids. 1mw8 or 1cy6 and such 280aa sequence had no homology to *M. tuberculosis* topoisomerase I sequence (protein id. CKN53907). While *E. coli* topo3a (protein id. BAA15551; 653aa; PDB id. 1I7D) is only 28% similarity at the NH_2_ terminus of human topo3a (protein id. ADP90493; 1001aa; PDB id. 4CGY) and no similarity to human topo3b implying NU2 likely is not an inhibitor of human topoisomerases. Similarly, very poor similarity of human nuclear and mitochondrial topo1B (nuclear protein id. AAA61207 and mitochondrial protein id. NP_001245375; 765aa and 601aa) to topo1A of *E. coli* (Protein id. BAA14811; 865aa). Note that *E. coli* topo1A omega protein carboxy terminus 273aa was not located in 67kd active topo1A of *E. coli* and crystallography was done with NH_2_ terminus 592aa (PDB id.1MW8).

Molecular modelling also showed some promise (figure-11). Keeping that promise MDR-cure lotion with five herbs mixture or each alone cured the MDR nail infections in three human patients and 83% lowered *E. coli* KT-1_mdr infections in rats. Still, more clinical research necessary to use NU2 and CU1 in human similar to antibiotics. However, quite pure CU1 and NU2 of *Cassia fistula* bark and *Suregada multiflora* root may be used in human to cure MDR infections similar to Ayurvedic drugs where clinical studies relaxed to some extent. The exposure of NU2 on Molly fishes and HeLa cells were not found at moderate concentration (figure-4). Our impression is that at high dose (>100ug/ml) NU2 should be toxic to animal and human where many topoisomerases could be affected.

## Conclusion

We have identified fluoropolyphosphate phytodrug NU2 from *Suregada multiflora* and such phytochemical active against MDR bacteria targeting topoI and topoIII and must be studied further in clinical trials in animal and then in human. We think antibiotic void will continue due to few reasons: (1) *mdr* genes are accumulated in large conjugative plasmids as well as chromosome islands with many transposons; (2) the spread of *mdr* genes in plasmids is increasing at 5%/year and (3) a critical message has generated to protect symbiosis relation between gut bacteria and human facilitating efficient *mdr* genes *(bla, aac, aph, aad*) creation to protect gut microbiota from antibiotics for continuous vitamin synthesis. Hence, new phyto-drug development is crucial to society and equally justified for developed countries like USA where every year more than 20000 deaths have been reported. In India, such deaths are increasing day by day. We have shown MDR-Cure extracts cures human MDR nail and skin infection. Importantly, taxol, artimisinin, reserpine, epipodophyllotoxin and quinine are well approved phytomedicines. We demonstrated that NU2 (as well as CU1) may be similar phyto-drugs with type-I DNA topoisomerase and or RNA polymerase targets. We do not know targets of CU3, JU2, NU3 and VU2 abundant phytochemicals with strong antibacterial activities. Such ancient Hindu Civilization method of herbal therapy against MDR bacteria as well as other unicellular parasites may save million people those are dragged into poverty line due to high cost of drugs and extended time hospitalization in poor countries.

## Competent interests

The author declared no conflict of interest.

## Ethical issues

The author declared no ethical issues as no human subject or animal was used in the study.

